# On the sparsity of fitness functions and implications for learning

**DOI:** 10.1101/2021.05.24.445506

**Authors:** David H. Brookes, Amirali Aghazadeh, Jennifer Listgarten

## Abstract

Fitness functions map biological sequences to a scalar property of interest. Accurate estimation of these functions yields biological insight and sets the foundation for model-based sequence design. However, the amount of fitness data available to learn these functions is typically small relative to the large combinatorial space of sequences; characterizing how much data is needed for accurate estimation remains an open problem. There is a growing body of evidence demonstrating that empirical fitness functions display substantial sparsity when represented in terms of epistatic interactions. Moreover, the theory of Compressed Sensing provides scaling laws for the number of samples required to exactly recover a sparse function. Motivated by these results, we develop a framework to study the sparsity of fitness functions sampled from a generalization of the NK model, a widely-used random field model of fitness functions. In particular, we present results that allow us to test the effect of the Generalized NK (GNK) model’s interpretable parameters—sequence length, alphabet size, and assumed interactions between sequence positions—on the sparsity of fitness functions sampled from the model and, consequently, the number of measurements required to exactly recover these functions. We validate our framework by demonstrating that GNK models with parameters set according to structural considerations can be used to accurately approximate the number of samples required to recover two empirical protein fitness functions and an RNA fitness function. In addition, we show that these GNK models identify important higher-order epistatic interactions in the empirical fitness functions using only structural information.

## Introduction

Advances in high-throughput experimental technologies now allow for the probing of the fitness of thousands, and sometimes even millions, of biological sequences. However, these measurements generally represent only a tiny fraction of those required to comprehensively characterize a fitness function. It is therefore critical to develop methods that can estimate fitness functions from an incomplete set of measurements. Many methods have been proposed for this purpose, ranging from the fitting of regularized linear models [1], and parameterized biophysical models [2, 3] to nonparametric techniques [4, 5], and various nonlinear machine learning approaches [6], including deep neural networks [7, 8]. In addition to providing basic biological insight, such methods have been used to improve the efficiency and success rate of experimental protein engineering approaches [9, 10, 11] and are crucial components of *in-silico* sequence design tools [12, 13, 14, 15].

Despite these advances in fitness function estimation, the answer to one fundamental question remains elusive—namely how many experimental fitness measurements are required to accurately estimate a fitness function. We refer to this problem as that of determining the sample complexity of fitness function estimation. Insights on this topic can be used to inform researchers on which of a variety of experimental techniques should be used to probe a particular fitness function of interest, and on how to restrict the scope of an experimental probe such that the resulting data allows one to accurately estimate the function under study. Our central focus herein is to elucidate the open question of the sample complexity of fitness function estimation.

It has recently been observed that some empirical fitness functions—those for which experimental fitness measurements are available for all possible sequences—are sparse when represented in the Walsh-Hadamard basis, which represents fitness functions in terms of all possible “epistatic” interactions (i. e., nonlinear contributions to fitness due to interacting sequence positions) [16, 17, 3]. Further, this sparsity property has been exploited to improve estimators of such functions [18, 19, 20]. Indeed, it is well known in the field of signal processing that sparsity enables more statistically efficient estimation of functions. Additionally, results from Compressed Sensing (CS), a sub-field of signal processing, provide scaling laws for the number of measurements required to recover a function in terms of its sparsity [21, 22]. These results suggest that by studying the sparsity of fitness functions in more depth, we may be able to predict the sample complexity of fitness function estimation.

Although an increasing number of empirical fitness functions are available that could allow us to investigate sparsity in particular example systems, these data necessarily only report on short sequences in limited environments. A common approach in evolutionary biology to overcome the lack of sufficient empirical fitness functions is to instead study ‘random field’ models of fitness, which assign fitness values to sequences based on stochastic processes constructed to mimic the statistical properties of natural fitness functions [23, 24]. We follow a similar line of reasoning and study the sparsity of fitness functions sampled from random field models, allowing us to probe properties of a much broader class of fitness functions than the available empirical data. We make use of a particular random field model, namely a generalization of the widely-used NK model [25]. The NK model is known to represent a rich variety of realistic fitness functions despite requiring only two parameters to be defined: *L*, the sequence length,^1^ and *K*, the maximum degree of epistatic interactions. In the NK model, each sequence position interacts with a “neighborhood” of *K* – 1 other positions that either include directly adjacent positions or are chosen uniformly at randomly [23]. NK models have been shown to model a number of properties of empirical fitness functions, including fitness correlation functions [26, 27] and adaptive walk statistics [25, 28, 29]. The Generalized NK (GNK) model [30], extends the model by allowing neighborhoods to be of arbitrary size and content. We refer to simulated fitness functions sampled from the GNK model as ‘GNK fitness functions’.

Buzas and Dinitz [30] calculated the sparsity of GNK fitness functions represented in the WH basis as a function of the sequence length and the composition of the neighborhoods. Nowak and Krug [31] expanded on this work by calculating the sparsity of GNK fitness functions with a few specific neighborhood schemes, as a function of only the size of the neighborhoods. Notably, these works consider only binary sequences, and use sparsity as a tool to understand the properties of adaptive walks on GNK landscapes, without connecting it to fitness function estimation. In contrast, our aim is to determine the sample complexity of estimating GNK fitness functions in the biologically-relevant scenarios where sequences are made up of non-binary elements (e.g., nucleotide or amino acid alphabets). In order to do so, we extend the results of refs. 30 and 31 to the case of non-binary alphabets by employing “Fourier” bases, which are generalizations of the WH basis that can be constructed for any alphabet size. We then leverage CS theory to determine the minimum number of measurements required to recover GNK fitness functions in the Fourier basis. This framework of using CS theory in tandem with the GNK model allows us to test the effects of sequence length, alphabet size, and interaction structure on the sample complexity of estimating GNK fitness functions.

We validate the practical utility of our framework by demonstrating that suitably parametrized GNK models can accurately approximate the sparsity of several empirical landscapes, and thus we can successfully leverage our sample complexity results to determine how many measurements are needed to estimate these landscapes. In particular, we use GNK models that incorporate structural information to show this for two empirical protein landscapes, and one ‘quasi-empirical’ RNA landscape. Our analysis also demonstrates that structure-based GNK models correctly identify many of the important higher-order epistatic interactions in the corresponding empirical fitness functions despite using only second-order structural contact information. This surprising insight bolsters a growing understanding of the importance of structural contacts in shaping fitness functions.

In the next sections, we summarize the relevant background material required for our main results.

### Fitness functions and estimation

A fitness function maps sequences to a scalar property of interest, such as catalytic efficiency [17], binding affinity [2], or fluorescent brightness [32]. In particular, let 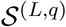 be the set of all *q^L^* possible sequences of length *L* whose elements are members of an alphabet of size *q* (e. g., *q* = 4 for nucleotides, and 20 for amino acids); then a fitness function is any function that maps the sequence space to scalar values, 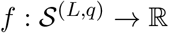. In practice, sequences may contain different alphabets at different positions, but these can usually be mapped to a common alphabet. For instance, one position in a nucleotide sequence may be restricted to A or T, and another to G or C, but both of these can be mapped onto the binary alphabet {0,1}. In the SI we consider the case where the size of the alphabet may be different at each position.

Any fitness function of sequences of length *L* and alphabet size *q* can be represented exactly as

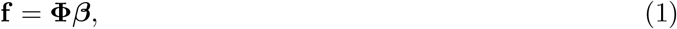

where **f** is the vector of all *q^L^* fitness values, one for each possible sequence, **Φ** is a *q^L^* × *q^L^* orthogonal basis, and ***β*** is the vector of *q^L^* coefficients corresponding to the fitness function in that basis. Although any orthogonal basis may be used, here we restrict **Φ** to refer to either the Walsh-Hadamard basis (when *q* = 2), or the Fourier basis (for *q* > 2), which will be defined shortly. Each row of **Φ** represents an encoding of a particular sequence in 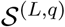. Now suppose we observe *N* fitness measurements, 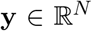, for *N* different sequences, each corresponding to one of the rows of **Φ**. The goal of fitness function estimation is then to recover a good approximation to ***β*** using these *N* measurements, which correspond to only a subset of all possible sequences. In general, this is an underdetermined linear system that requires additional information to be solved, and many methods have been developed for this purpose. The extent to which a fitness function is recovered by such a method can be assessed by the mean squared error (MSE) between the estimated and true coefficients. Since **Φ** is an orthogonal matrix, this is equivalent to the MSE between the true fitness values **f** and those predicted using the estimated coefficients.

The field of Compressed Sensing is primarily concerned with studying algorithms that can solve underdetermined systems, and specifying the conditions under which recovery with a specified amount of error in the estimated coefficients can be guaranteed. Therefore, it stands to reason that CS may be helpful for characterizing fitness function estimation problems. The LASSO algorithm is among the most widely-used and well-studied for solving underdetermined systems, both in CS and also in machine learning [33]. The key determinant of success of algorithms such as LASSO in recovering a particular function is how sparse that function is when represented in a particular basis, or how well it can be approximated by a function that is sparse in that basis. Using the fitness function estimation problem as an example, a central result from CS [34] states that if ***β*** is an *S*-sparse vector (meaning that is has exactly *S* nonzero elements), then with high probability LASSO can recover ***β*** exactly with

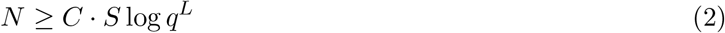

noiseless fitness measurements, where *C* is an unknown constant. For this bound to hold, the *N* sequences with observed fitness measurements must be sampled uniformly from the space of sequences [34]. It has also been shown that if ***β*** is only approximately sparse (i. e., it has many small, but nonzero, coefficients), or if there is noise in the measurements, then the error in the recovery can still be bounded (Materials and Methods).

Eq. 2 shows that if we are able to calculate the sparsity of a fitness function, and estimate a value for the constant *C*, then we can calculate the number of samples required to recover that fitness function with LASSO. Note that the “sparsity” of a fitness function is defined as the number of nonzero coefficients when the fitness function is expanded in a particular basis.^2^ Sparsity is defined with respect to the particular orthonormal basis which must therefore be chosen carefully. In the next section, we discuss bases that can be used to represent fitness functions.

### Fourier bases for fitness functions

The sparsity of a class of natural signals depends crucially on the basis with which they are represented. Many fitness functions have been shown to be sparse in the Walsh-Hadamard (WH) basis [16, 17, 3], which has also been used extensively in theoretical studies of fitness landscapes [35, 27, 36, 37] and even to unify multiple definitions of epistasis [38]. The WH basis can be interpreted as encoding fitness functions in terms of epistatic interactions [39, 38]. In particular, when a fitness function of binary sequences of length *L* is represented in the form of Eq. 1 (with **Φ** being the WH basis), then the sequence elements are encoded as {–1, 1} and the fitness function evaluated on a sequence, **s** = [*s*_1_, *s*_2_, …, *s_L_*], has the form of an intuitive multi-linear polynomial [20],

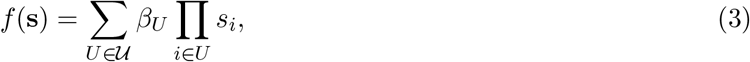

where 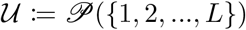 is the power set of sequence position indices. Each of the 2^*L*^ elements of 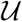 is a set of indices that corresponds to a particular epistatic interaction, with the size of that set indicating the order of the interaction (e. g., if a 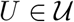 is of size |*U*| = *r*, then it represents an *r*^th^ order interaction). The coefficient *β_U_* is an element of ***β***, indexed by its corresponding epistatic interaction set.

The WH basis can only be used to represent fitness functions of binary sequences, which poses a challenge in biological contexts where common alphabets include the nucleotide (*q* = 4) and amino acid (*q* = 20) alphabets. This issue is typically skirted by encoding elements of a larger alphabet as binary sequences (e. g., by using a “one-hot encoding”), and using the WH basis to represent fitness functions of these encoded sequences. However, doing so results in an inefficient representation, which has dramatic consequences on the calculation of sample complexities. To see this, consider the “one-hot” encoding scheme of amino acids, where each amino acid is encoded as a length 20 bit string. The number of amino acid sequences of length *L* is 20^*L*^, while the one-hot encodings of these sequences are elements of a binary sequence space of size 2^20*L*^ = 1,048, 576^*L*^. This latter number also corresponds to the number of WH coefficients required to represent the fitness function in the one-hot encoding, and is much too large to be of any practical use.

Although it is not widely recognized in the fitness function literature, it is possible to construct bases analogous to the WH basis for arbitrarily-sized alphabets, which we refer to as “Fourier” bases (Materials and Methods, 40, 41). The WH basis is the Fourier basis for *q* = 2. The Fourier basis for a larger alphabet shares much of the WH basis’s intuition of encoding epistatic interactions between positions in a sequence. In particular, we have an analogous expression to Eq. 3 for the Fourier basis, in which the fitness function is represented as a sum of 2^*L*^ terms, each of which corresponds to an epistatic interaction. In the WH basis, an *r*^th^ epistatic interaction *U* in a sequence *s* is encoded as the scalar 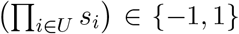, while in the Fourier basis, it is represented by a length (*q* – 1)^*r*^ vector, which we denote as ***ϕ**_U_*(**s**). Similarly, in the WH basis each epistatic interaction is associated with a single coefficient, while in the Fourier basis, each epistatic interaction is associated with (*q* – 1)^*r*^ coefficients. All together, the evaluation of a fitness function represented in the Fourier basis on a sequence **s** is given by

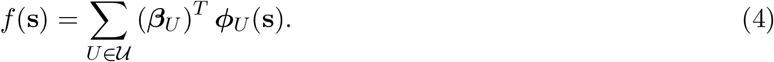

It is shown below that when GNK fitness functions are represented in the Fourier basis, then we have the intuitively pleasing result that all of the Fourier coefficients associated with a particular epistatic interaction are identically distributed, and thus the GNK model can be interpreted in terms of epistatic interactions.

### The Generalized NK model

Sampling fitness functions from a random field model provides a means to simulate fitness functions of sequences of any length or alphabet size. A random field model specifies a stochastic process that assigns fitness values to all possible sequences. This process implicitly defines a joint probability distribution over the fitness values of all sequences, and another over all of the Fourier coefficients, ***β***.

Herein, we focus on the Generalized NK (GNK) model [30]. In order to be defined, the GNK model requires the specification of the sequence length *L*, alphabet size *q*, and an interaction “neighborhood” for each position in the sequence. A neighborhood, *V*^[*j*]^, for sequence position *j* is a set of position indices that contains *j* itself, and *K_j_* – 1 other indices, where we define *K_j_* ≔ |*V*^[*j*]^| to be the size of the neighborhood. Given *L*, *q*, and a neighborhood for each position, the GNK model assigns fitness to every sequence in the sequence space via a series of stochastic steps (Materials and Methods). In the GNK model, two sequences have correlated fitness values to the extent that they share subsequences corresponding to the positions in the neighborhoods. For example, consider a GNK model defined for nucleotide sequences of length 3, where the first neighborhood is *V*^[1]^ = {1, 3}. Then the sequences ACG and AAG will have partially correlated fitness values because they both contain the subsequence AG in positions 1 and 3. One of the key intuitions of the GNK model is that larger neighborhoods will produce more “rugged” fitness functions in which many fitness values are uncorrelated, because it is less likely for two sequences to share subsequences when the neighborhoods are large. Note that larger neighbourhoods also implies the presence higher order epistatic interactions.

The key choice in specifying the GNK model is in how the neighborhoods are constructed. We will characterize the sparsity induced by three “standard” schemes for constructing neighborhoods [31, 36]: the Random, Adjacent and Block Neighborhood schemes. These schemes all restrict every neighborhood to be the same size, *K*, which provides a basis for comparing how different interaction structures induce sparsity in fitness functions. Graphical depictions of these three schemes are shown in Fig. 1. We will additionally consider a novel scheme where neighborhoods are constructed based on contacts between residues in an atomistic protein structure, which is described in more detail in a later section.

**Fig. 1:**
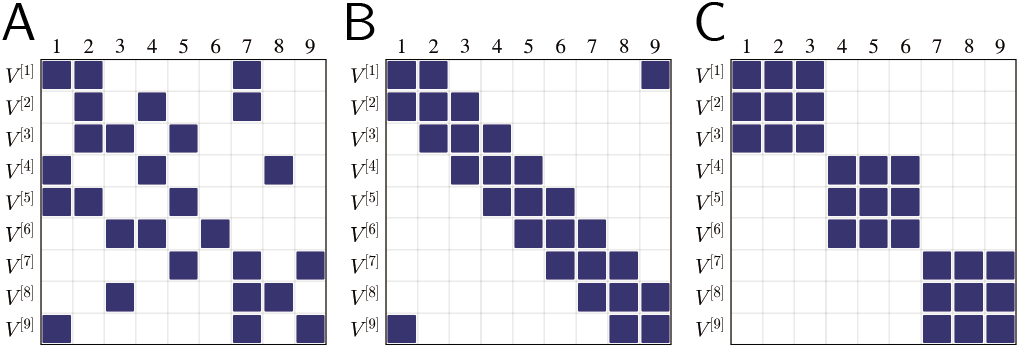
Graphical depictions of GNK neighborhood schemes for *L* = 9 and *K* = 3. Rows on the vertical axis represent neighborhoods and columns on the horizontal axis represents sequence positions. A square in the (*i*, *j*)^th^ position in the grid denotes that sequence position *j* is in the neighborhood *V*^[*i*]^. (a) Random Neighborhoods (b) Adjacent Neighborhoods (c) Block Neighborhoods.

Notably, the GNK model is an example of a spin glass, a popular model in statistical physics, with different neighborhood schemes corresponding to different types of spin glasses [42]. Further, the recovery of sparse spin glass Hamiltonians has been investigated in some depth [43].

In the next section, we present results that enable us to calculate the sparsity of GNK fitness functions given the sequence length, alphabet size, and a set of neighborhoods. The proofs of these results are given in the SI.

## Results

### The sparsity of GNK fitness functions

A somewhat remarkable feature of the GNK model is that it can be shown that the Fourier coefficients of GNK fitness functions are independent normal random variables whose mean and variance can be calculated exactly given the sequence length, alphabet size, and neighborhoods. In particular, the Fourier coefficients of fitness functions sampled from the GNK model are distributed according to 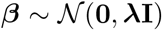, where **λ** is a vector of variances corresponding to each element of ***β*** and **I** is the *q^L^* × *q^L^* identity matrix. Further, each of the (*q* – 1)^*r*^ Fourier coefficients corresponding to an *r*^th^ order epistatic interaction, *U*, have equal variances given by

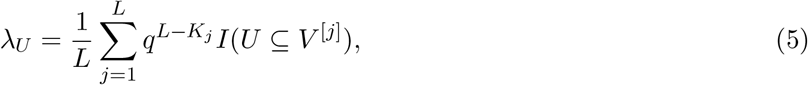

where, with a slight abuse of notation, *I*(*U* ⊆ *V*^[*j*]^) is an indicator function that is equal to one if *U* is a subset of the neighborhood *V*^[*j*]^, and zero otherwise. Eq. 5 shows that the variance of a Fourier coefficient is roughly proportional to the number of neighborhoods that contain the corresponding epistatic interaction as a subset. Most importantly for our purposes, Eq. 5 implies that a Fourier coefficient only has nonzero variance when the corresponding epistatic interaction is a subset of at least one neighborhood; otherwise the coefficients are deterministically zero. Consequently, we can use Eq. 5 to calculate the total number of Fourier coefficients that are not deterministically zero in a specified GNK model, which is equal to the sparsity of all fitness functions sampled from the model. In particular, the sparsity, *S*(*f*), of a fitness functions *f* sampled from a GNK model is given by

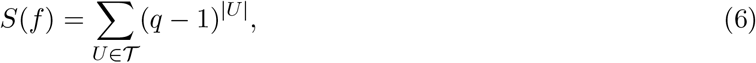

where 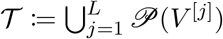 is the union of the powersets of the neighborhoods. Eq. 6 makes the connection between neighborhoods and epistatic interactions concrete: the GNK model assigns nonzero Fourier coefficients to any epistatic interactions whose positions are included in at least one of the neighborhoods. For example, if positions 3 and 4 in a sequence are both in some neighborhood *V*^[*j*]^, then all elements of ***β***_{3, 4}_ are nonzero. Further, by the same reasoning, the coefficients corresponding to all subsets of positions {3, 4} are also nonzero (i. e., the coefficients corresponding to the first order effects associated with positions 3 and 4).

Eq. 6 provides a general formula for the sparsity of GNK fitness functions as a function of *L*, *q* and the neighborhoods. We can use this formula to calculate the sparsity of GNK fitness functions with each of the standard neighborhood schemes—Random, Adjacent and Block—for a given neighborhood size, *K*. In the Materials and Methods, we provide exact results for the sparsity of GNK fitness functions with Adjacent and Block neighborhoods, and the expected sparsity of GNK fitness functions with for Random neighborhoods. We also provide an upper bound on sparsity of GNK fitness functions with any neighborhood scheme with constant neighborhood size, *K*. In Figs. 2A and 2B we plot this upper bound for a variety of settings of *L*, *q* and *K*. Further, in Fig. 2C we plot the upper bound along with the exact or expected sparsity of GNK fitness functions with each of the standard neighborhood schemes. We can see that even at the same setting of *K*, different neighborhood schemes result in striking differences in the sparsity of sampled fitness functions.

**Fig. 2:**
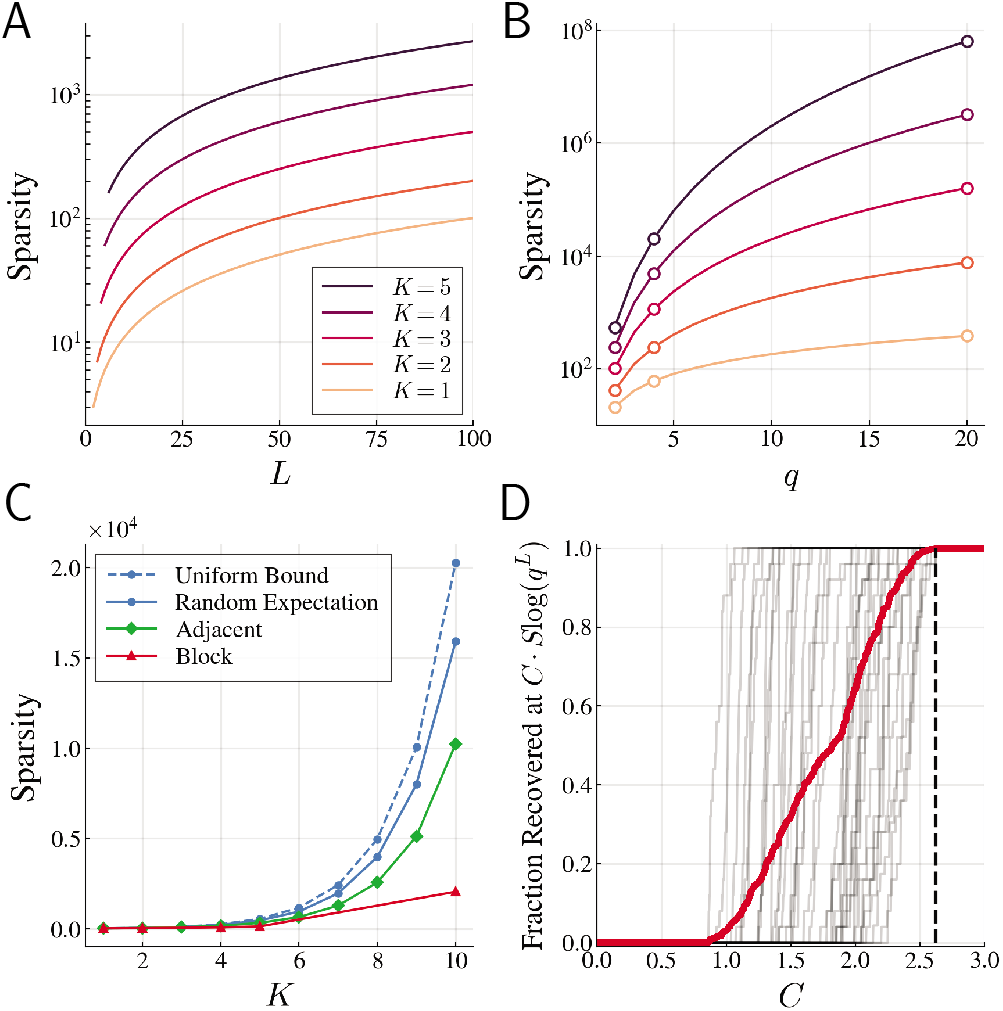
The sparsity of GNK fitness functions. (A) Upper bound on sparsity of GNK fitness functions with constant neighborhoodmeasurements required to exactly recover sizes for *q* = 2 and a range of settings of the *L* and *K* parameters (B) Upper bound for *L* = 20 and a range of settings of the alphabet size *q* and the *K* parameter (colors as in (A)). Alphabet sizes corresponding to binary (*q* = 2), nucleotide (*q* = 4), and amino acid (*q* = 20) alphabets are highlighted with open circles. (C) Sparsity of GNK fitness functions with neighborhoods constructed with each of the standard neighborhood schemes for *L* = 20 and *q* = 2, and and a range of settings of *K*, denoted by markers. (D) Fraction of sampled GNK fitness functions with Random Neighborhoods recovered at a range of settings of *C*. Each grey curve represents sampled fitness functions at a particular values of *L* ∈ {5, 6, …, 13}, *q* ∈ {2, 3, 4} and *K* ∈ {1, 2, 3, 4, 5}. The red curve averages over all 907 sampled functions. The value *C* = 2.62, which we chose to use for subsequent numerical experiments, is highlighted with a dashed line.

### Exact recovery of GNK fitness functions

The sparsity result of Eq. 6 allows us to apply CS theory to determine the number of fitness measurements required to recover GNK fitness functions exactly. Specifically, we can use Eq. 2 to determine a minimal *N* such that exact recovery is guaranteed for an *S*(*f*)-sparse fitness function *f* when there is no measurement noise. However, to do so, we first need to determine an appropriate value for the constant *C* in Eq. 2, which we did via straightforward numerical experiments. In particular, we used LASSO to estimate GNK fitness functions using varying numbers of randomly sampled, noiseless fitness measurements and from these estimates, determined the minimum number of training samples required to exactly recover the fitness functions (allowing for a small amount of numerical error–see Materials and Methods for more details). We then determined the minimum value of *C* such that Eq. 2 holds in each tested case. Fig. 2D summarizes the experiments, showing that *C* = 2.62 is sufficiently large to ensure recovery of all of the over 900 tested fitness functions, and we use this value for all further calculations. A more detailed analysis of these experiments is shown in Fig. S3, which makes clear that the minimum possible setting of *C* is a function of *L*, *q*, and *K*, and therefore that *C* = 2.62 may be a conservative setting for certain reasonable settings of these parameters.

We next used this estimate of *C*, along with our results for the sparsity of GNK fitness functions, and the CS result of Eq. 2, to determine the minimum number of measurements required to exactly recover GNK fitness functions. Figs. 3A and 3B show examples of these calculations, where we used the bound on sparsity for GNK fitness functions with constant neighborhood sizes to calculate an upper bound on the minimum number of samples required to recover these fitness functions. A number of important insights can be derived from Fig. 3. First, the number of measurements required to perfectly estimate these fitness functions is many orders of magnitude smaller than the total size of sequence space. Consider, for instance the point in Fig. 3a where *L* = 50 and thus the size of sequence space is 2^50^ ≈ 10^15^, 10 orders of magnitude greater than the largest plotted sample complexity. Additionally, comparing Figs. 3A and 3B clearly indicates that increasing the alphabet size within biologically relevant ranges increases the number of samples required to recover fitness functions at a faster rate than increasing the length of the sequence.

**Fig. 3:**
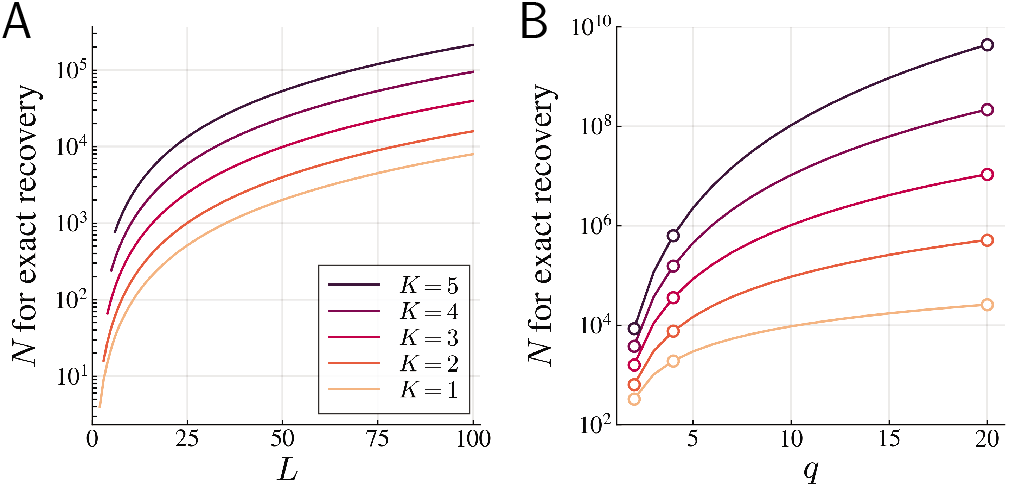
Minimum number of measurements required to exactly recover GNK fitness functions with constant neighborhood sizes. (A) Upper bound on the minimum *N* required to recover GNK fitness functions with constant neighborhood sizes for *q* = 2 and a range of settings of the *L* and *K* parameters. (B) Upper bound for *L* = 20 and a range of settings of the alphabet size *q* and the *K* parameter (colors as in (A)). Alphabet sizes corresponding to binary (*q* = 2), nucleotide (*q* = 4), and amino acid (*q* = 20) alphabets are highlighted with open circles.

### Analysis of empirical protein fitness functions

In order to validate our framework, we next tested the extent to which our results could be used to predict the sample complexity of estimating empirical protein fitness functions. To do so, we made use of a novel scheme for constructing GNK neighborhoods, which we call the Structural neighborhood scheme, that uses information derived from 3D structure of a given protein. In particular, Structural neighborhoods are constructed based on contacts between amino acid residues in a given atomistic protein structure where, following refs. 4 and 45, we define two residues to be in contact if any two atoms in the residues are within 4.5Å of each other. Then the Structural neighborhood of a position *j* contains all positions that are in structural contact with it.

An interesting aspect of the Structural neighborhood scheme is how it encodes epistatic interactions through Eq. 6. In particular, in a GNK model with Structural neighborhoods, higher-order epistatic interactions arise from only pairwise structural contact information—that is, an *r*^th^ order epistatic interaction has nonzero Fourier coefficients when *r* – 1 positions are in structural contact with a central position..

We instantiated GNK models with Structural neighbourhoods for two proteins: the TagBFP fluorescent protein (PDB: 3M24 [46]) and the protein encoded by the His3 gene in *Saccharomyces cerevisiae* (His3p). We then used the results described in the previous section to calculate the sparsity of GNK fitness functions with these Structural neighborhoods, the sample complexity of estimating these functions, and the variance of each the functions’ Fourier coefficients.

Both TagBFP and His3p are associated with empirical fitness functions with complete or nearly complete sets of experimental measurements. We calculated the Fourier coefficients associated with each of these empirical fitness functions using Ordinary Least Squares (or regression with a small amount of regularization when the measurements were only nearly complete), so as to be able to compare the resulting sparsity and magnitude of the empirical Fourier coefficients to those of the corresponding GNK fitness functions with Structural neighborhoods. Next, to assess whether the sample complexity of estimating GNK fitness functions with Structural neighborhoods can be used to inform the sample complexity of estimating real protein fitness functions, we fit LASSO estimates of the empirical fitness functions with varying numbers of randomly sampled empirical measurements, and determined how well each recovered the empirical fitness function.

In the case of the TagBFP structure, the associated empirical fitness function contains functional observations (blue fluorescence brightness) of mutations to the mTagBFP2 protein [18], which is closely related to TagBFP but has no available structure. This data contains measurements for all combinations of mutations in 13 positions, where each position is allowed to mutate to only one other amino acid (i.e., *L* = 13 and *q* = 2), yielding 2^13^ = 8192 total fitness observations. A graphical depiction of the Structural neighborhoods associated with these 13 positions is shown in the first row of Fig. 4A. Using Eq. 6 for the GNK model with these Structural neighborhoods yielded a sparsity of *S*(*f*) = 56, while application of Eq. 5 enabled us to determine the distribution of these 56 non-zero Fourier coefficients and the epistatic interactions that they corresponded to.

**Fig. 4:**
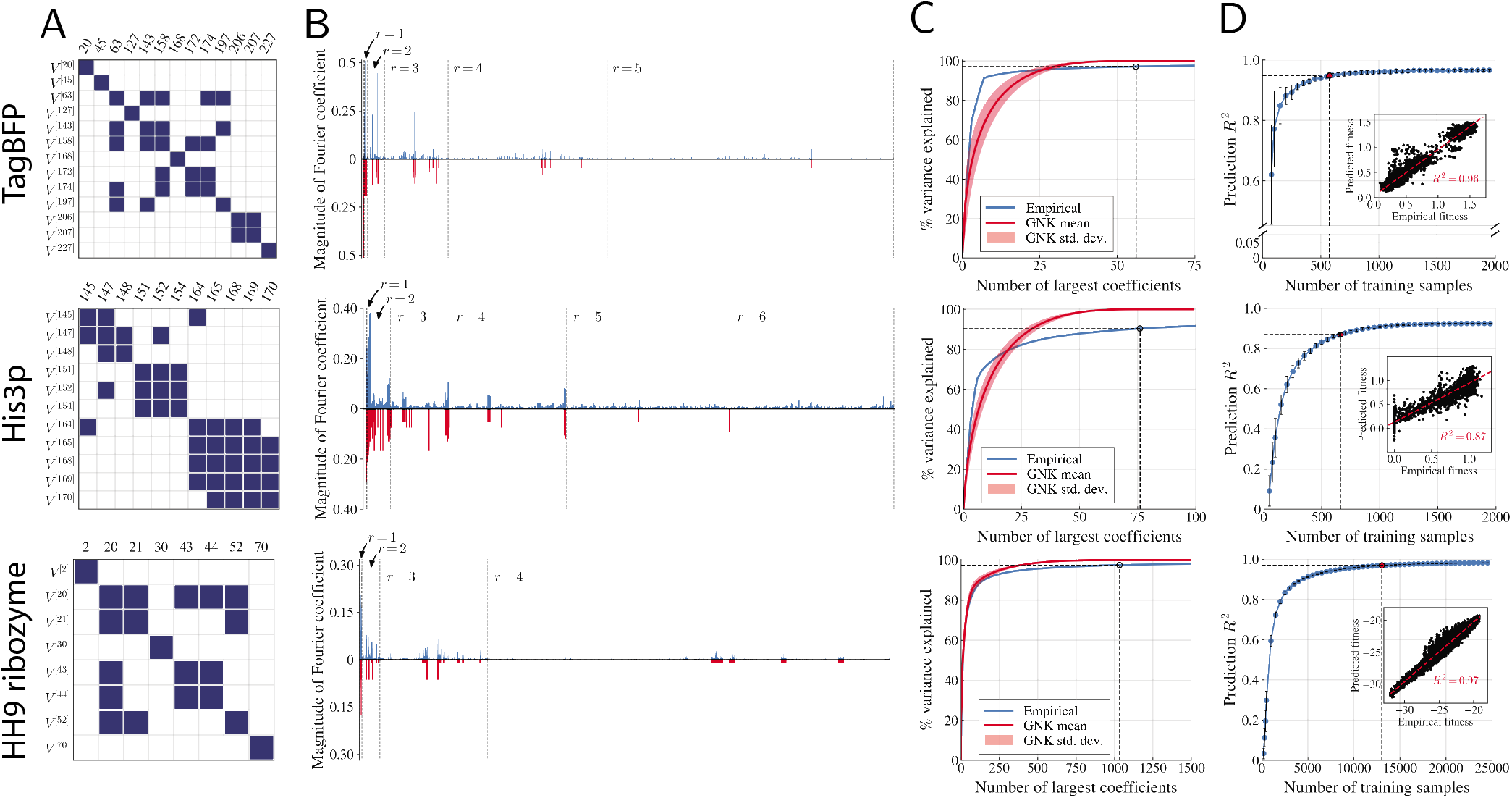
Comparison of empirical fitness functions to GNK models with Structural Neighborhoods. *First row*: comparison to mTagBFP2 fitness function from ref. 18. *Second row*: comparison to His3p fitness function from ref. 44. *Third row*: comparison to quasi-empirical fitness function of the Hammerhead ribozyme HH9. (A) Structural Neighborhoods derived from crystal structural of TagBFP (first row), I-TASSER predicted structure of His3p (second row), and predicted secondary structures of the Hammerhead Ribozyme HH9 (third row). (B) Magnitude of empirical Fourier coefficients (upper plot, in blue) compared to expected magnitudes of coefficients in the GNK model (reverse plot, in red). Dashed lines separate orders of epistatic interactions, with each group of *r*^th^ order interactions indicated. (C) Percent variance explained by the largest Fourier coefficients in the empirical fitness functions and in fitness functions sampled from the GNK model. The dotted line indicates the exact sparsity of the GNK fitness functions, which is 56 is in the first row 76 in the second, and 1,033 in the third, at which points 97.1%, 90.4%, and 97.5% of the empirical variances are explained, respectively. (D) Error of LASSO estimates of empirical fitness functions at a range of training set sizes. Each point on the horizontal axis represents the number of training samples, *N*, that are used to fit the LASSO estimate of the fitness function. Each point on the blue curve represents the *R*^2^ between the estimated and empirical fitness functions, averaged over 50 randomly sampled training sets of size *N*. The point at the number of samples required to exactly recover the GNK model with Structural Neighborhoods (*N* = 575 in the first row, *N* = 660 in the second, and *N* = 13, 036 in the third) is highlighted with a red dot and dashed lines; at this number of samples, the mean prediction *R*^2^ is 0.948 in the first row, 0.870 in the second, and 0.969 in the third. Error bars indicate the standard deviation of *R*^2^ over training replicates. Insets show paired plots between the estimated and predicted fitness function for one example training set of size *N* = 575 (first row), *N* = 660 (second row), and *N* = 13, 036 (third row).

For the case of His3p, we used a nearly combinatorial complete empirical fitness function that is embedded in the data of ref. 44. In particular, the data contains 2030 out of the possible 2048 fitness observations for sequences corresponding to 11 positions in His3p, each taking on one of two amino acids (i. e., *L* = 11 and *q* = 2). We constructed Structural neighborhoods based on the I-TASSER [47] predicted structure of His3p [44] (Fig. 4A, second row), which resulted in sparsity *S*(*f*) = 76 for GNK fitness functions with these neighborhoods, and we again computed the distribution of these coefficients and determined the corresponding epistatic interactions.

The comparisons of the mTagBFP and His3p empirical fitness functions with the associated GNK models with Structural neighborhoods are summarized in Fig. 4. First, we examined the magnitudes of the Fourier coefficients of the empirical and GNK fitness functions. Since the Fourier coefficients in the GNK model are independent normal random variables, the expected magnitude of a coefficient with variance λ is 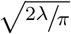. A comparison of all coefficients corresponding to up to 5^th^ and 6^th^ order epistatic interactions are shown in Fig. 4B for the mTagBFP and His3p cases, respectively. Many of the epistatic interactions with the largest empirical coefficients also have nonzero coefficients in the GNK model with Structural neighborhoods, suggesting that these models are reasonable approximations to protein fitness functions. In the SI, we quantify the overlap between the largest coefficients in the empirical and GNK fitness functions by performing statistical tests that show that the coefficients identified as being nonzero by the GNK model have significantly higher ranks in the empirical coefficients than those identified as being zero (Figs S10–S13)

Although none of the empirical Fourier coefficients are exactly zero, these coefficients display substantial approximate sparsity. In particular, over 95% and 80% of the total variance in the coefficients can be explained by the 25 coefficients with the largest magnitude in the mTagBFP and His3p fitness functions, respectively. To more holistically assess whether GNK fitness functions with Structural neighborhoods approximate the sparsity of the empirical fitness functions well, we compared the percent variance explained by the *S* Fourier coefficients with the largest magnitudes in both the empirical and GNK fitness functions, for a range of settings of *S*. Fig. 4C shows the results of this comparison, with the blue curve showing the percent variance explained by the largest empirical coefficients, and the red curve and red shaded region showing the mean and standard deviation, respectively, of the percent variance explained by the largest coefficients in 1,000 sampled GNK fitness functions. Considering that these plots show only the first few of all possible coefficients that could be included on the horizontal axis (75 out of the 8,192 for mTagBFP and 100 out of 2,048 for His3p), it is clear that the GNK model approximates the sparsity of the empirical fitness function qualitatively well. Of particular importance is the point at which all of the nonzero coefficients of the GNK fitness functions are included in the calculation (i.e., 100% of the variance is explained), which occurs at *S* = 56 and 76 in the mTagBFP and His3p cases, respectively; at this point, more than 90% of the empirical variance is explained in both cases.

These promising sparsity comparisons suggest that the sample complexity of estimating GNK fitness functions with Structural neighborhoods may be used to approximate the number of measurements required to effectively estimate protein fitness functions. We confirmed this by using LASSO to estimate the empirical fitness functions with varying number of training points and regularization parameter chosen by cross-validation (Fig. 4D). Our theory predicts that 548 and 630 samples are minimally needed for exact recovery of the GNK fitness functions with mTagBFP and His3p Structural neighborhoods, respectively. In both cases, we see these sample sizes produce effective estimates of the corresponding empirical fitness functions, with a mean *R*^2^ of 0.95 and 0.87 for estimates of the mTagBFP and His3p fitness functions, respectively.

In the SI we show analogous results to those in Fig. 4 for another nearly complete subset of the His3p fitness data of ref. 44 that contains 48,219 out of 55,296 fitness measurements for the same 11 positions discussed above and alphabets that differ in size at each position. Altogether, these results suggest the GNK model with Structural neighborhoods can be used to approximate the sparsity of protein fitness functions, and the sample complexity of estimating such functions.

### Analysis of a quasi-empirical RNA fitness function

As further validation, we next tested the ability of our framework to predict the sample complexity of estimating a quasi-empirical RNA landscape. In particular, we studied the fitness function of all possible mutations to the *Erinaceus Europaes* Hammerhead ribozyme HH9 wild type sequence (RFAM: AANN01066007.1) at positions 2, 20, 21, 30, 43, 44, 52, and 70 where the fitness of each sequence in this *L* = 8, *q* = 4 sequence space is given by the Minimum Free Energy (MFE) of the secondary structures associated with the sequence, as calculated by the ViennaRNA package [48]. We follow [49] in referring to this as a ‘quasi-empirical’ fitness function, as it is constructed from an established physical model rather than direct experimental measurements. The magnitudes of the Fourier coefficients associated with this fitness function are shown as blue bars in the third row of Fig. 4B. This is a sparse landscape, with the largest 150 out of 65,536 possible coefficients explaining over 90% of the quasi-empirical variance.

We then used a GNK model with RNA-specific Structural neighborhoods to predict the sample complexity of estimating this quasi-empirical landscape. In order to construct these neighborhoods, we first used ViennaRNA to sample 10,000 secondary structures from the Boltzmann ensemble of structures associated with the wild-type sequence. We then built neighborhoods where a position *j* was included in the neighborhood of position *k* if (i) *j* and *k* were directly adjacent in the sequence or (ii) *j* and *k* were paired in any of the sampled secondary structures (Fig. 4A, third row). The expected magnitude of the Fourier coefficients in the GNK model with these neighborhoods are shown as red bars in Fig. 4B. Once again we see that the GNK model with Structural neighborhoods identifies many of the most important higher-order epistatic interactions in this fitness function.

As with the empirical protein fitness functions, we compared the sparsity of the GNK and quasi-empirical fitness functions (Fig. 4C, third row) and tested the ability of our framework to predict the sample complexity of estimating the quasi-empirical fitness function with LASSO (Fig. 4D, third row). These results demonstrate that a suitably parameterized GNK model can accurately model the sparsity of a realistic RNA fitness function, which bolsters our results on empirical protein fitness functions and further suggests that the GNK model can be a practical tool for estimating the sample complexity of fitness function estimation.

## Discussion

By leveraging perspectives from the fields of Compressed Sensing and evolutionary biology, we developed a framework for calculating the sparsity of fitness functions and the number of fitness measurements required to exactly recover those functions with the LASSO algorithm (or another sparse recovery algorithm with CS guarantees) under a well-defined set of assumptions. These assumptions are that (i) the fitness functions are sampled from a specified GNK model, (ii) fitness measurements are noiseless, (iii) fitness measurements correspond to sequences sampled uniformly at random from the space of sequences, and (iv) the fitness functions are represented in the Fourier basis. Under these assumptions, our results allow us to test the effect of sequence length, alphabet size, and positional interaction structure on the sparsity and sample complexity of fitness function estimation.

We have additionally demonstrated that in certain cases our results can be used to estimate the sample complexity of estimating protein fitness functions when assumptions (i) and (ii) may not be exactly satisfied. In particular, we showed that GNK models with Structural neighborhoods accurately approximate the sparsity of two empirical protein fitness functions and a quasi-empirical RNA fitness function, and can be used to estimate the number of measurements required to recover those empirical fitness functions with high accuracy. The success of applying our framework to these fitness functions, which are neither exactly sparse nor noiseless (in the case of the protein fitness functions), is at least partially due to the fact that sparse recovery algorithms such as LASSO are robust to approximate sparsity and noisy measurements (Materials and Methods, Eq. 8).

It should be noted that assumptions (iii) and (iv) likely result in conservative estimates for the sample complexity of fitness function estimation. Uniform sampling of sequences is optimal when one has no *a priori* knowledge about the fitness function; however, if one knows which coefficients in a fitness function are likely to be nonzero, then it may be possible to construct alternative sampling schemes, or deterministic sets of sequences to measure, such that the fitness function can be recovered with many fewer measurements than with uniform sampling. Additionally, it may be possible to construct a basis in which certain classes of fitness functions are more sparse than in the Fourier basis, and this will in turn result in fewer measurements being required to recover those fitness functions when they are represented in the alternative basis.

Our sample complexity predictions could be used to a certain extent to guide experimental probes of fitness by suggesting how one should restrict the scope of mutagenesis such that recovery of the resulting fitness function can be expected with a certain amount of data. Using protein mutagenesis experiments as an example, this could be done by limiting the number of positions that are mutated, perhaps based on biophysical considerations [50, 3] or previous experimental results [51, 52, 53, 54], or by allowing each position to mutate to only a restricted alphabets of amino acids, for instance by choosing only amino acids that are present in homologous sequences [18, 44, 17]. Of course, one should take care not to minimize the sample size requirements at the expense of probing important areas of the protein or nucleotides sequences under study.

Complementing our main contributions, we have also demonstrated that GNK models with Structural neighborhoods can predict the identity of many of the largest higher-order epistatic interactions in empirical protein fitness functions (Fig. 4B, first and second rows). There are a number of false positives (i.e. coefficients that the GNK model identifies as nonzero, but are very small in the empirical fitness function) and false negatives in these plots that deserve some comment. To explain these errors, it is first important to remember that the red bars in Fig. 4B represent the *expected* magnitudes of zero-mean GNK coefficients; even among fitness functions sampled directly from the GNK model, we would expect to see “false positives” where the sampled magnitudes were smaller than the expected magnitudes. The false negatives may be explained by three similar causes, all regarding the insufficiency of using a single crystal or predicted structure to construct Structural neighborhoods for proteins. First, the structures we used may simply be inaccurate: in one case, we use the TagBFP crystal structure, while the fitness function reports on mutations to mTagBFP2; in the His3p case we use an I-TASSER predicted structure that may have inaccuracies. Secondly, static structures do not capture dynamical effects that may impact fitness; for instance two residues may be in contact in a non-native conformation of the protein that differs from the crystallized or predicted conformation. Finally, the crystal or predicted structures of wild-type proteins cannot capture the potential structural changes that may occur when the protein is mutated, as is done to collect fitness data. Additionally, we used a fixed contact threshold of 4.5Å, but adjusting this threshold can moderately change the GNK Fourier coefficients (Figs. S4–S7); most notably the largest empirical *r* = 6 coefficient in the His3p fitness function is identified as being nonzero by the GNK model when we increase the cutoff distance to 7Å.

Few attempts have been made at understanding how many measurements are required to estimate fitness functions, despite the practical importance of this question for experimental design. By making the connection between this question and the known sparsity of fitness functions in certain bases, we provide a much-needed framework for probing the sample complexity of estimating fitness functions. Further, we show that the GNK model, given protein and RNA structural information, can gauge the sparsity of empirical fitness functions enough to make useful statements about the sample complexity of estimating such functions. As data collection progresses, the tools and understanding to probe sample complexity may have to correspondingly progress, but our work provides a solid foundation on which to do so.

## Materials and Methods

### Compressed Sensing

As described in the main text, the fitness function estimation problem is to solve the underdetermined linear system **y** = **X*β*** for an unknown ***β***, where **y** is a vector of *N* fitness measurements, and **X** is a matrix containing the *N* corresponding rows of **Φ** that represent the sequences with fitness measurements. Herein we assume that each element of **y** is corrupted with independent Gaussian noise with variance *σ*^2^. LASSO solves for an estimate of the Fourier coefficients by solving following convex optimization program:

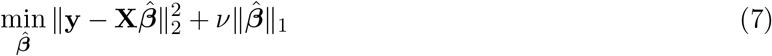

where *ν* is a hyperparameter that determines the strength of regularization. Candes and Plan [34] proved that when the rows of an orthogonal basis such as **Φ** are sampled uniformly at random, and the number of samples satisfies Eq. 2, then the solution to the program in Eq. 7, denoted ***β***\*, satisfies with high probability

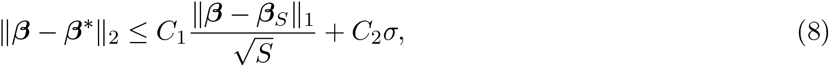

where *C*_1_, and *C*_2_ are constants and ***β***_*S*_ is the best *S*-sparse approximation to ***β***, i.e., the vector that contains the *S* elements of ***β*** with the largest magnitude and sets all others elements to zero. Eq. 8 has a number of important implications. First, it tells us that if ***β*** is itself *S*-sparse, then, in a noiseless setting, it can be recovered *exactly* with 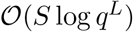 measurements. Otherwise, if ***β*** is not exactly sparse but is well approximated by a sparse vector, then it can be approximately recovered with error on the order of 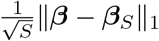, which is proportional to the sum of the magnitudes of the *q^L^* – *S* elements of ***β*** with the smallest magnitudes.

We primarily focus on cases where a fitness function is exactly sparse in the Fourier basis and we can calculate the sparsity. Although natural fitness functions are unlikely to be exactly sparse, they may be well approximated by sparse vectors, and Eq. 8 tells us that the error of the estimator will be well controlled in this case. Similarly, measurement noise in experimental fitness data is unavoidable, but Eq. 8 shows that the error induced by this noise is dependent on the variance of the measurement noise, and not on the properties of the fitness function itself. Since here we are primarily concerned with understanding how assumed properties of fitness functions affect the sample complexity of estimating those functions, it is thus most appropriate to consider the noiseless setting and leave the estimation of error due to measurement noise to future work.

### Fourier bases

Our generalization of the WH basis to larger alphabets is based on the theory of Graph Fourier bases. The Graph Fourier basis corresponding to a given graph is a complete set of orthogonal eigenvectors of the Graph Laplacian of the graph. Graph Fourier bases have many useful properties and have been used extensively for processing signals defined on graphs [55].

The WH basis is specifically the Graph Fourier basis corresponding to the Hamming graph *H*(*L*, 2) [56]. The vertices of *H*(*L*, 2) represent all unique binary sequences of length *L*; two sequences are adjacent in *H*(*L*, 2) if they differ in exactly one position (i.e., the Hamming distance between the two sequences is equal to one). The Hamming graphs *H*(*L*, *q*) are defined in the same way for sequences with alphabet size *q*. Thus, we can construct an analogous Graph Fourier basis to the WH basis to represent sequences with larger alphabets by calculating the eigenvectors of the Graph Laplacian of *H*(*L*, *q*). Since we only consider functions defined on Hamming graphs, we refer to Graph Fourier bases corresponding to Hamming graphs simply as Fourier bases.

An important property of the Hamming graph *H*(*L*, *q*) is that it can be constructed as the *L*-fold Graph Cartesian product of the “complete graph” of size *q* [56]. The complete graph of size *q*, denoted *K*(*q*), has *q* vertices (which represent elements of the alphabet in our case) and edges between all pairs of vertices. Due to the spectral properties of graph products, the eigenvectors of the Hamming graph (i. e., the Fourier basis) can be calculated as a function of the eigenvectors of the complete graph. An orthonormal set of eigenvectors of the Graph Laplacian of the complete graph *K*(*q*) is given by the columns of the following Householder matrix:

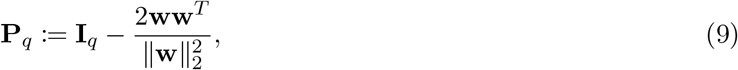

where 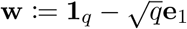, **1**_*q*_ is the vector of length *q* whose elements are all equal to one, **e**_1_ is the length *q* with the first element set to 1 and all others set to zero, and **I***_q_* is the *q* × *q* identity matrix.

The complete graph is equal to the Hamming graph *H*(1, *q*), and thus Equation Eq. 9 constructs the Fourier basis for sequences of length one and alphabet size *q*. Each row of **P**_*q*_ corresponds to a sequence of length one; the first column is constant for all rows while the remaining *q* – 1 columns encode the alphabet elements (i.e., the final *q* – 1 elements of a row uniquely identify the alphabet element that the row corresponds to). More specifically, let 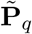 be the matrix containing the final *q* – 1 unnormalized columns of **P**_*q*_, such that 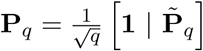, where | denotes column-wise concatenation. Then the *i*^th^ row of 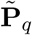 encodes the *i*^th^ element of the alphabet; we denote each of these encodings as **p**_*q*_(*s*), where *s* is an element of the alphabet (i. e., each **p**_*q*_(*s*) is a row of 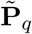).

Then, it can be shown that the Fourier basis corresponding to the Hamming graph *H*(*L, q*), which can be used to represent fitness functions of sequences of length *L* and alphabet size *q*, is given by the *L*-fold Kronecker product of the eigenvectors of the complete graph. More concretely, an orthonormal set of eigenvectors of the Graph Laplacian of the Hamming graph *H*(*L*, *q*) is given by the columns of following the *q^L^* × *q^L^* matrix [57]:

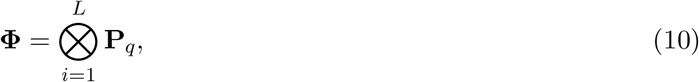

where **P**_*q*_ is defined in Eq. 9. In the basis defined in Eq. 10, an epistatic interaction between positions in the set *U* is encoded by the length (*q* – 1)^|*U*|^ vector 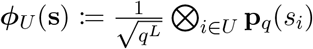. These encodings are used in the Fourier basis representation of fitness functions shown in Eq. 4. The results of Eq. 9 and Eq. 10 are proved in the SI. Note that an equivalent form of this basis for *q* = 4 was given in ref. 40 and an alternative form for any alphabet size was given in ref. 41.

### GNK Model

Given sequence length, *L*, alphabet size, *q*, and set of neighborhoods 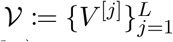, a fitness function sampled from the GNK model assigns a fitness to every sequence 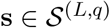 with the following two steps:

1. Let **s**^[*j*]^ ≔ (*s_k_*)_*k*∈*V*^[*j*]^_ be the subsequence of **s** corresponding to the indices in the neighborhood *V*^[*j*]^. Assign a ‘subsequence fitness’, *f_j_*(**s**^[*j*]^) to every possible subsequence, **s**^[*j*]^, by drawing a value from the normal distribution with mean equal to zero and variance equal to ^1^/*L*. In other words, 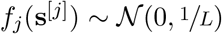 for every 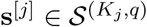, and for every *j* = 1, 2, …, *L*.
2. For every 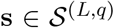, the subsequence fitness values are summed to produce the total fitness values 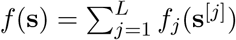.

This definition of the GNK model is slightly more restrictive than that presented in ref. 30. In particular, in ref. 30 the authors allow subsequence fitness values to be sampled from any appropriate distribution whereas for simplicity we consider only the case where subsequence fitness values are sampled from the scaled unit normal distribution.

### Standard neighborhood schemes

We consider three standard neighborhood schemes: the Random, Adjacent and Block neighborhood schemes. In all of these, each neighborhood is of the same size, *K* (i. e., *K_j_* = *K* for all *j* = 1, 2, …, *L*). In the Random scheme, each neighborhood *V*^[*j*]^ contains *j* and *K* – 1 other position indices selected uniformly at random from {1, 2, …, *L*}\*j*. In the Adjacent scheme when *K* is an odd number, each neighborhood *V*^[*j*]^ contains the 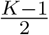 positions immediately clockwise and counterclockwise to *j* when the positions are arranged in a circle. When *K* is an even number, the neighborhood includes the 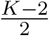 counterclockwise positions and the 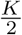 clockwise positions. The Block scheme (also known as the Block Model [58, 59]), splits positions into 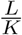 blocks of size *K* and lets each block be “fully connected” in the sense that every neighborhood of a position in the block contains all other positions in the block, but no positions outside of the block. In order for Block neighborhoods to be defined, *L* must be a multiple of *K*.

### Standard neighborhood sparsity calculations

The sparsity of GNK fitness functions with the standard neighborhood schemes can be calculated exactly as functions of *L*, *q*, and *K*. The following results are used in the main text and are all proved in the SI. First, the sparsity of any GNK fitness with uniform neighborhood sizes is bounded above by

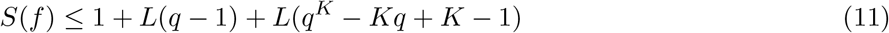

All curves in Figs. 2A and 2B are calculated with this bound, and it is also used for the sample complexity calculations shown in Fig. 3. It is also plotted as the dashed blue curve in Fig. 2C with *L* = 20 and *q* = 2. Additionally, the sparsity of GNK fitness functions with Block neighborhoods can be calculated exactly and is given by

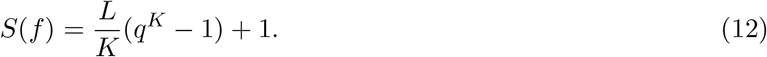

Eq. 12 is plotted as the red curve in Fig. 2C with *L* = 20 and *q* = 2. Similarly, the sparsity of GNK fitness functions with Adjacent neighborhoods is given by

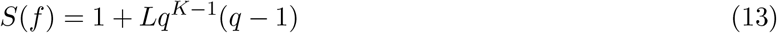

which is plotted as the green curve in Fig. 2C with *L* = 20 and *q* = 2. Finally, the expected sparsity of GNK fitness functions with Random neighborhoods, with the expectation taken over the randomly assigned neighborhoods, is given by

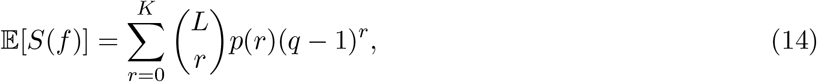

where

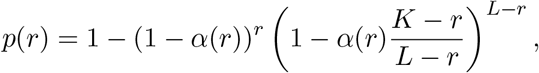

and 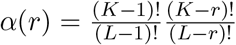. Eq. 14 with *L* = 20 and *q* = 2 is shown as the solid blue curve in Fig. 2C. The results of Eqs. **-14** are proved in the SI.

### Numerical calculation of *C*

In order to determine an appropriate value of *C*, we (i) sampled a fitness function from a GNK model, (ii) subsampled *N* sequence-fitness pairs uniformly at random from the complete fitness function for a range of settings of *N*, (iii) ran LASSO on each of the subsampled data sets and (iv) determined the smallest *N* such that the fitness function is exactly recovered by LASSO. Letting 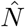 be the minimum *N* for which exact recovery occurs, then

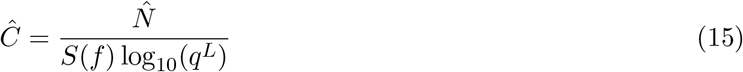

is the minimum value of *C* that satisfies Eq. 2, where *S*(*f*) is calculated with Eq. 6. We ran multiple replicates of this experiment for neighborhoods sampled according to the RN scheme, for different settings of *L*, *q* and *K*. This resulted in a test for 907 total fitness functions. For each of these fitness functions, we ran LASSO with 5 randomly sampled training sets for each size *N*, and a regularization parameter, *ν*, determined by cross-validation. We deemed the fitness function exactly recovered when the estimates resulting from all 5 training sets explained 99.99% of the variance in the fitness function’s coefficients.

Equipped with an estimate of *C*, we can calculate the minimum number of samples required to exactly recover a GNK fitness function by using Eq. 2 along with the sparsity calculations discussed in the previous section. Specifically,

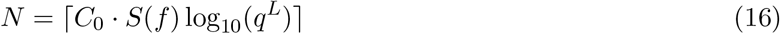

is the minimum number of samples that guarantees exact recovery, where ⌈⌉ represents the ceiling operator. Eq. 16 was used along with the bound in Eq. to calculate the curves in Fig. 3.

### Percent variance explained

In Fig. 4C, we computed the percent of total variance in the Fourier coefficients explained by the *S* coefficients with the largest magnitudes, for a range of settings of *S*. The percent variance explained by the *S* largest elements of the vector of coefficients ***β*** is calculated as

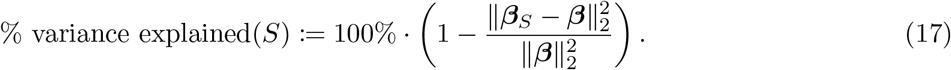

### Data and code availability

The data sets and code used for our analyses are available at https://github.com/dhbrookes/FitnessSparsity.

## Acknowledgements

We thank Akosua Busia and Chloe Hsu for helpful comments on the manuscript. D.H.B and J.L. were supported by the Chan Zuckerberg Investigator Program. A.A. was supported by the ARO (W911NF2110117).

## S1 Additional details on mTagBFP2 fitness function

The data of ref. 18 reports on the fluorescence of every intermediate between the mTagBFP2 blue fluorescent and mKate2 red fluorescent proteins. These two proteins differ in only 13 positions, and the red and blue fluorescence of all 2^13^ possible combinations of the two sequences at these positions was tested. Table S1 shows the alphabet of each position in these empirical fitness functions. For our analysis of this data, we considered only the reported blue fluorescence of each tested sequence. Since there is no available structure for mTagBFP2, we calculated the Structural neighborhoods with the structure of TagBFP, a blue fluorescent protein from which mTagBFP2 is derived by making the mutations S2_S2delinsVSKGE/I174A [60]. No crystal structure is available for mKate2 or a closely related protein (e. g., mKate) that would have allowed to us to perform analogous analysis on the red fluorescent data.

Since the empirical fitness function in the data of ref. 18 is combinatorially complete, we can solve for the Fourier coefficients with Ordinary Least Squares (OLS) regression. In particular, letting **y** be the empirical fitness values and **Φ** the WH basis encoding all binary sequences of length 13, then the OLS estimate of the empirical Fourier coefficients is 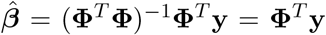, where the second equality is due to the orthogonality of **Φ**. The magnitudes of 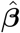 are plotted in Figure 4B (first row).

**Table S1:** Alphabets at each mutated position in the empirical fitness function of ref. 18. The first row indicates the index of the position in the complete protein sequence, the second row and third rows indicate the amino acids present at these positions in the mTagBFP2 and mKate2 sequences, respectively.

| Position | 20 | 45 | 63 | 127 | 143 | 158 | 168 | 172 | 174 | 197 | 206 | 207 | 231 |
| --- | --- | --- | --- | --- | --- | --- | --- | --- | --- | --- | --- | --- | --- |
| mTagBFP2 | D | V | L | T | F | N | S | A | A | Y | N | N | K |
| mKate2 | N | A | M | P | S | A | G | C | L | R | D | K | R |

## S2 Additional details and results on His3p fitness function

The complete data of ref. 44 reports the fitness of 875,151 unique amino acid sequences of the protein encoded by the His3 gene in yeast (which we refer to as His3p). In this case, the fitness was defined as the cellular growth rate when the the sequences were expressed in a strain of yeast. Embedded within this data, there exist a number of nearly combinatorially complete fitness functions. By ‘embedded’ we mean that if one only considers data reporting on a subset of the mutated positions, and a subset of the possible mutations at those positions, then nearly all combinations of the considered mutations at the considered positions have fitness values associated with them. Two such nearly complete fitness functions can be constructed by considering the 11 sequences positions 145, 147, 148, 151, 152, 154, 164, 165, 168, 169, and 170. If one considers only the two most frequently occurring amino acids at these positions, then 2,030 out of the 2^11^ = 2,048 possible combinations of those amino acids at those positions have fitness data associated with them. The two most frequent amino acids at each of these positions are shown in Table S2. We will refer to the fitness function corresponding to these 11 positions and alphabets as the His3p(small) fitness function. The His3p(small) empirical fitness function is analyzed in the main text and compared to GNK fitnesss functions with Structural neighborhoods (Figure 4, second row). These Structural neighborhoods were constructed using the I-TASSER predicted structure of His3p that is analyzed in ref. 44. Determining the empirical Fourier coefficients for the His3p(small)) fitness function requires solving a very slightly underdetermined linear system (2,030 rows and 2,048 columns). In order to do so, we solved for a LASSO estimate using all 2,030 fitness measurements with a small amount of regularization (*ν* = 1 × 10^−12^); the magnitudes of the resulting coefficients are shown in the second row of Figure 4B.

If one further considers larger alphabets at certain positions, then there is another nearly complete empirical fitness function associated with the 11 positions that are considered in the His3p(small) fitness function. In particular, consider alphabets of size **q** = [2, 2, 3, 2, 2, 3, 3, 4, 2, 4] corresponding to each of the 11 positions; the particular alphabets corresponding to each position are shown in Table S3. We refer to the fitness function corresponding to these alphabets as the His3p(big) fitness function. The data of ref. 44 contains fitness measurements for 48,219 out of the 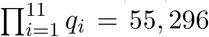 sequences in the space corresponding to the alphabets in Table S3. We show how to construct Fourier bases for fitness functions of sequences with hybrid alphabet sizes in Section S7. The Fourier basis for *L* = 11 sequence positions and hybrid alphabet sizes **q** has 55,296 associated coefficients. We used LASSO with *ν* = 1 × 10^−9^ to solve for the Fourier coefficients of the His3p(big) fitness functions. The magnitudes of these coefficients corresponding to up to 4^th^ order epistatic interactions are shown in blue in Figure S1A. We additionally show in Section S7 how to extend our results on GNK fitness functions to the case of hybrid alphabet sizes. We used these results to calculate the sparsity of GNK fitness functions with *L* = 11, alphabet sizes **q** and Structural Neighborhoods as in Figure 4D in the main text, as well as the the number of measurements required to recover those fitness functions. Figures S1A and S1B and show the results of comparing the Fourier coefficients and sparsity of the His3p(big) empirical fitness function to the GNK fitness functions (these are calculated with the same methodology as Figures 4B and 4C). Figure S1C shows the result of estimating the His3p(big) fitness function with randomly sampled measurements (analogous to Figure 4D in the main text). We again see that GNK fitness functions with Structural Neighborhoods can approximate the sparsity of, higher-order epistatic interactions in, and number of samples required to estimate empirical protein fitness functions. These results also demonstrate that the Fourier bases produce sparse representations of fitness functions with non-binary alphabets.

**Table S2:**
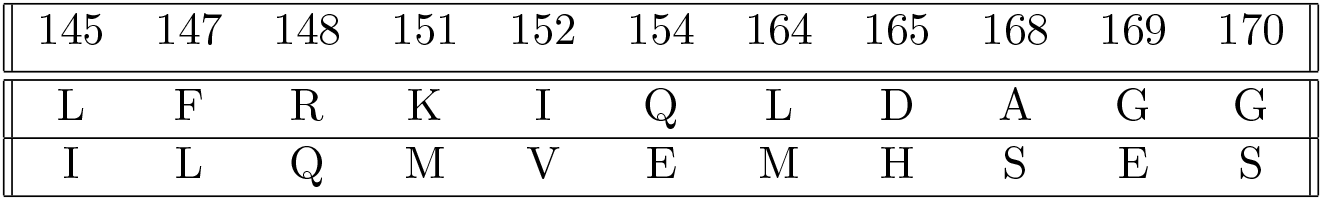
Alphabets at each mutated position in the His3p(small) fitness function. The first row indicates the sequence position. and the rows indicate amino acids that make up the alphabet at each position.

|  |  |  |  |  |  |  |  |  |  |  |
| --- | --- | --- | --- | --- | --- | --- | --- | --- | --- | --- |
| 145 | 147 | 148 | 151 | 152 | 154 | 164 | 165 | 168 | 169 | 170 |
| L | F | R | K | I | Q | L | D | A | G | G |
| I | L | Q | M | V | E | M | H | S | E | S |

**Table S3:** Alphabets at each mutated position in the His3p(big) fitness function. The first row indicates the sequence position. and the rows indicate amino acids that make up the alphabet at each position.

|  |  |  |  |  |  |  |  |  |  |  |
| --- | --- | --- | --- | --- | --- | --- | --- | --- | --- | --- |
| 145 | 147 | 148 | 151 | 152 | 154 | 164 | 165 | 168 | 169 | 170 |
| L | F | R | K | I | Q | L | D | A | G | G |
| I | L | Q | M | V | E | M | H | S | E | S |
|  |  | K |  |  | H | I | E |  | R | A |
|  |  |  |  |  | D |  | Q |  | Q | T |

**Fig. S1:**
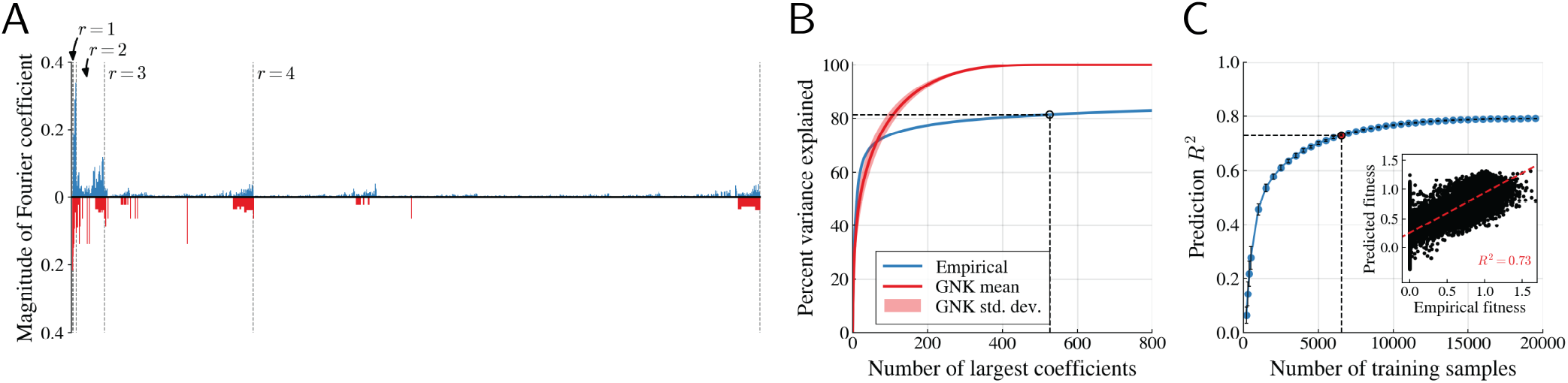
Comparison of His3p(big) empirical fitness function to the GNK model with Structural Neighborhoods from Figure 4A (second row) in the main text. (A) Magnitude of empirical Fourier coefficients (upper plot, in blue) compared to the standard deviations of coefficients in the GNK model (reverse plot, in red). Dashed lines separate orders of epistatic interactions, with each group of *r*^th^ order interactions indicated. (B) Percent of variance explained by the largest Fourier coefficients in the empirical fitness function and in fitness functions sampled from the GNK model. The dotted line indicates the exact sparsity of the GNK fitness functions (526) at which point 81.4% of the empirical variances is explained. (C) Error of LASSO estimates of empirical fitness functions at a range of training set sizes. Each point on the horizontal axis represents the number of training samples, *N*, that are used to fit the LASSO estimate of the fitness function. Each point on the blue curve represents the *R*^2^ between the estimated and empirical fitness functions, averaged over 50 randomly sampled training sets of size *N*. The point at the number of samples required to exactly recover the GNK model with Structural Neighborhoods (*N* = 6, 537) is highlighted with a red dot and dashed lines; at this number of samples, the mean prediction *R*^2^ is 0.728. Error bars indicate the standard deviation of *R*^2^ over training replicates. Inset shows paired plot between the estimated and predicted fitness function for one example training set of size *N* = 6, 537.

## S3 Additional details on quasi-empirical Hammerhead ribozyme HH9 fitness function

In order to construct the quasi-empirical fitness function of the *Erinaceus Europaes* Hammerhead ribozyme HH9 that is discussed in the main text, we first chose 8 positions on the wild type sequence of the ribozyme (RFAM: AANN01066007.1) to mutate. We chose positions by hand based on the predicted Minimum Free Energy (MFE) secondary structure of the wild type sequence, which is shown below in Fig. S2, with the aim of choosing positions that were at a range of distances from one another in the predicted structure. Ultimately we chose positions 2, 20, 21, 30, 43, 44, 52, and 70, and created the list of 4^8^ = 65, 536 sequences with all combinations of nucleotide substitutions at these positions. We then used the ViennaRNA package [48] to predict the Minimum Free Energy (MFE) of each sequence, which we used as the fitness value of each sequence. We then solved for the Fourier coefficients of this fitness function with OLS regression. In particular, letting **Φ** be the Fourier basis for sequences with *L* = 8 and *q* = 4, and **y** be the corresponding vector of MFE values, we estimate the Fourier coefficients, 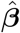 of the fitness function as 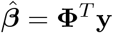. The magnitudes of 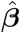 are shown as blue bars in the third row of Fig. 4B.

**Fig. S2:**
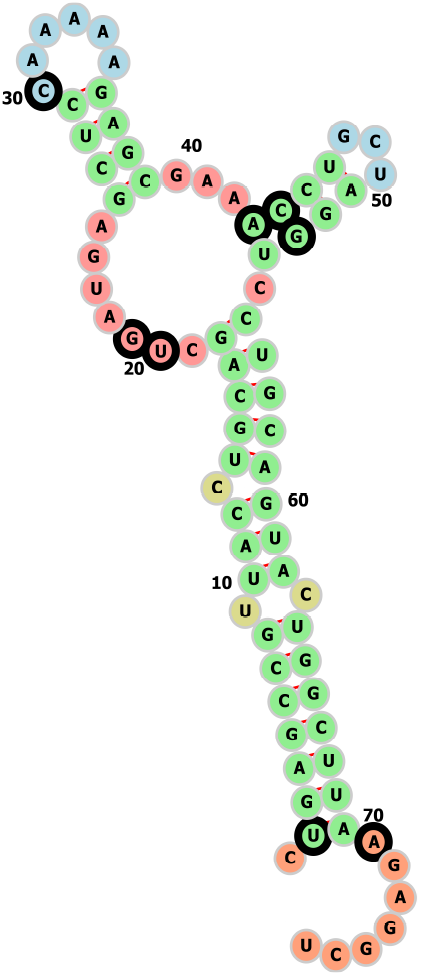
Minimum free energy secondary structure of of the *Erinaceus Europaes* Hammerhead ribozyme HH9 wild type sequence, as predicted by ViennaRNA [48] and visualized with forna [61]. The positions that were chosen to mutate are indicated with thick black outlines.

## S4 Additional details on numerical calculation of *C*

In order to determine an appropriate value of the *C* constant, we sampled fitness functions from the GNK model with Random Neighborhoods at different settings of *L*, *q*, and *K*, and determined the number of randomly sampled fitness measurements required to estimate these fitness functions. Figure 2D shows the results of these tests averaged over all settings of *L*, *q*, and *K*. In Figure S3, we display these results in more detail by showing the results for particular settings of *L* and *q* in separate plots. The curve in the plots of Figure S3 are also displayed as grey curves in Figure 2D.

**Fig. S3:**
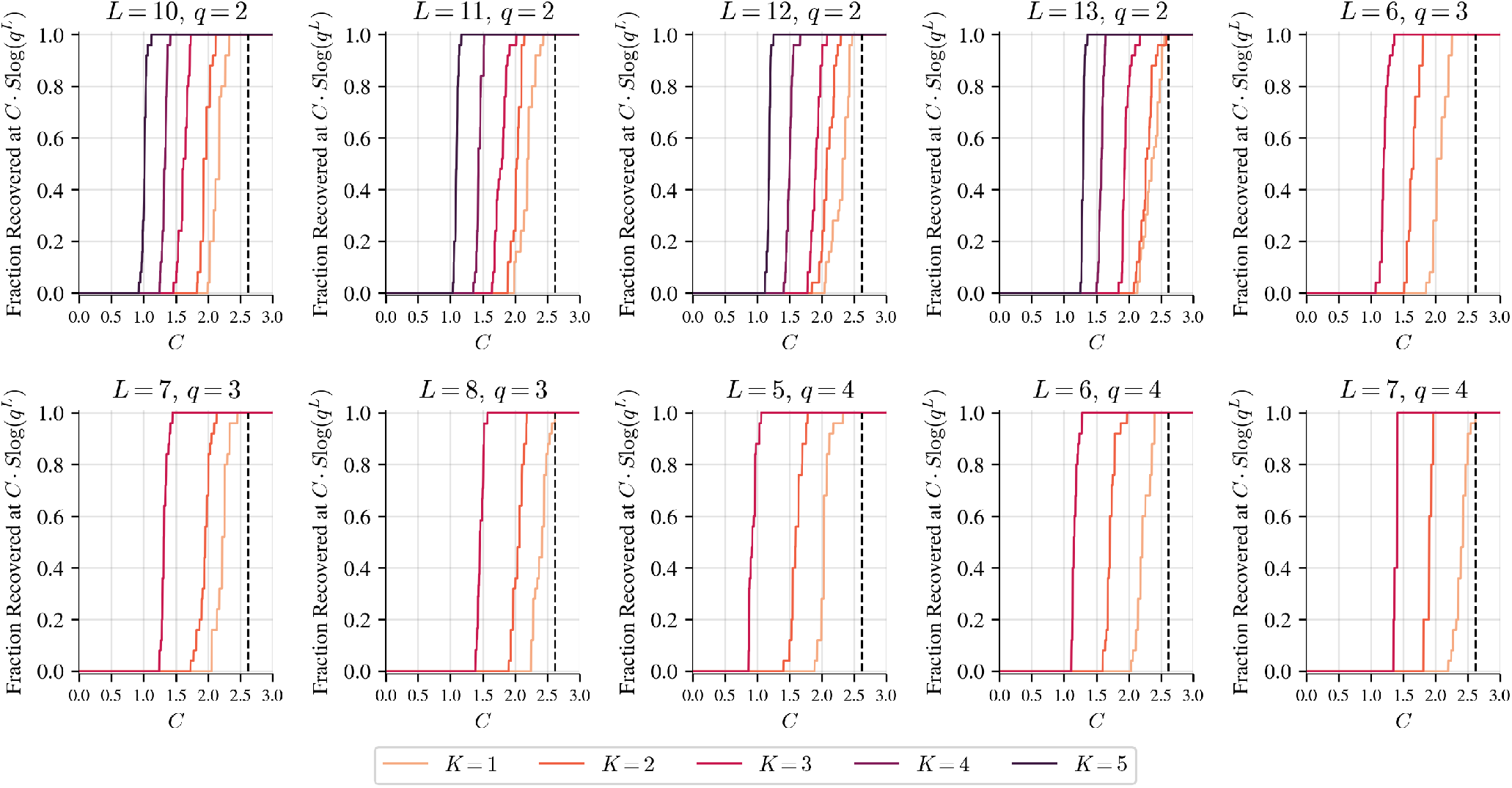
Fraction of GNK fitness functions with Random Neighborhoods recovered at a range of settings of *C*. Each plot corresponds to fitness functions with the setting of *L* and *q* indicated above the plot. Colors indicate the value of *K* used when sampling Random Neighborhoods. The value *C* = 2.62 is highlighted with a dashed line in each plot.

## S5 Additional analyses of empirical protein fitness functions

In this section, we present further analyses of the TagBFP and His3p fitness functions discussed in the main text and the associated GNK models with Structural neighborhoods.

### S5.1 Effect of contact threshold on Fourier coefficients of GNK fitness functions with Structural Neighborhoods

In the main text, we deem two positions of a protein sequence to be in structural contact when any pair of atoms in the residues at these positions are within a threshold distance of 4.5Å of one another in a given atomistic protein structure. We then used this definition of structural contacts to constructed Structural neighborhoods for GNK fitness functions. Here we test how modifying the threshold distance for defining structural contacts affects various properties of the Structural neighborhoods, and the GNK models that use these neighborhoods.

One interpretation of the Structural neighborhoods is as a binarization of the pairwise distance matrix between pairs of positions. In other words, given a pairwise distance matrix and a threshold distance, one can simply set distances less than the threshold distance equal to 1 and all other equal to zero to produce the graphical depictions of the Structural neighborhoods shown in the first and second rows of Fig. 4A. The non-binarized pairwise distance matrices of TagBFP and His3p are shown below in Fig. S4; these matrices provide more detail on the structural relationships between the positions and allows us to assess the effect of different thresholds on the composition of the Structural neighborhoods. By binarizing these distance matrices, one can construct the Structural neighborhoods corresponding to any threshold distance.

To assess the effect of modifying the threshold distance on GNK models with Structural neighborhoods, we determined the Structural neighborhoods corresponding to a range of threshold distances in both the TagBFP and His3p cases. We then Eq. Eq. 5 to calculate the variance of the Fourier coefficients of GNK fitness functions that used the Structural neighborhoods at each threshold distance. The magnitudes of these Fourier coefficients compared to the magnitudes of the corresponding empirical Fourier coefficients are shown in Figs. S5 (for the TagBFP fitness function) and S6 (for the His3p fitness function). We then used Eq. Eq. 2 to calculate the sparsity of GNK fitness functions with Structural neighborhoods constructed using a range of threshold distances; the results of these calculations are shown in S7.

**Fig. S4:**
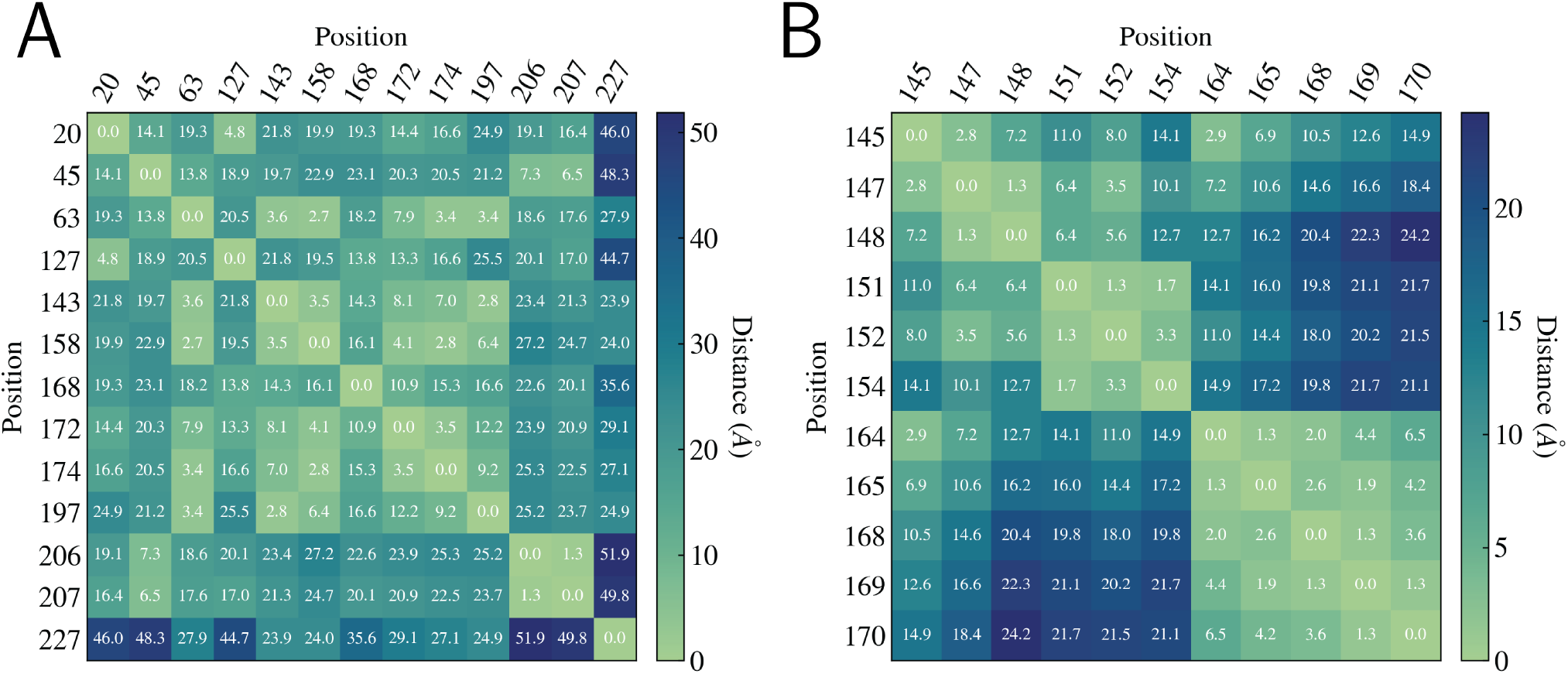
Pairwise distance matrices of the TagBFP (A) and His3p (B) structures, for positions that are mutated in the corresponding fitness functions. Each value in the grid reports the minimum distance between any pair of atoms in the residues at the positions indicated in the labels of the grid. The grids are colored based on these distances.

**Fig. S5:**
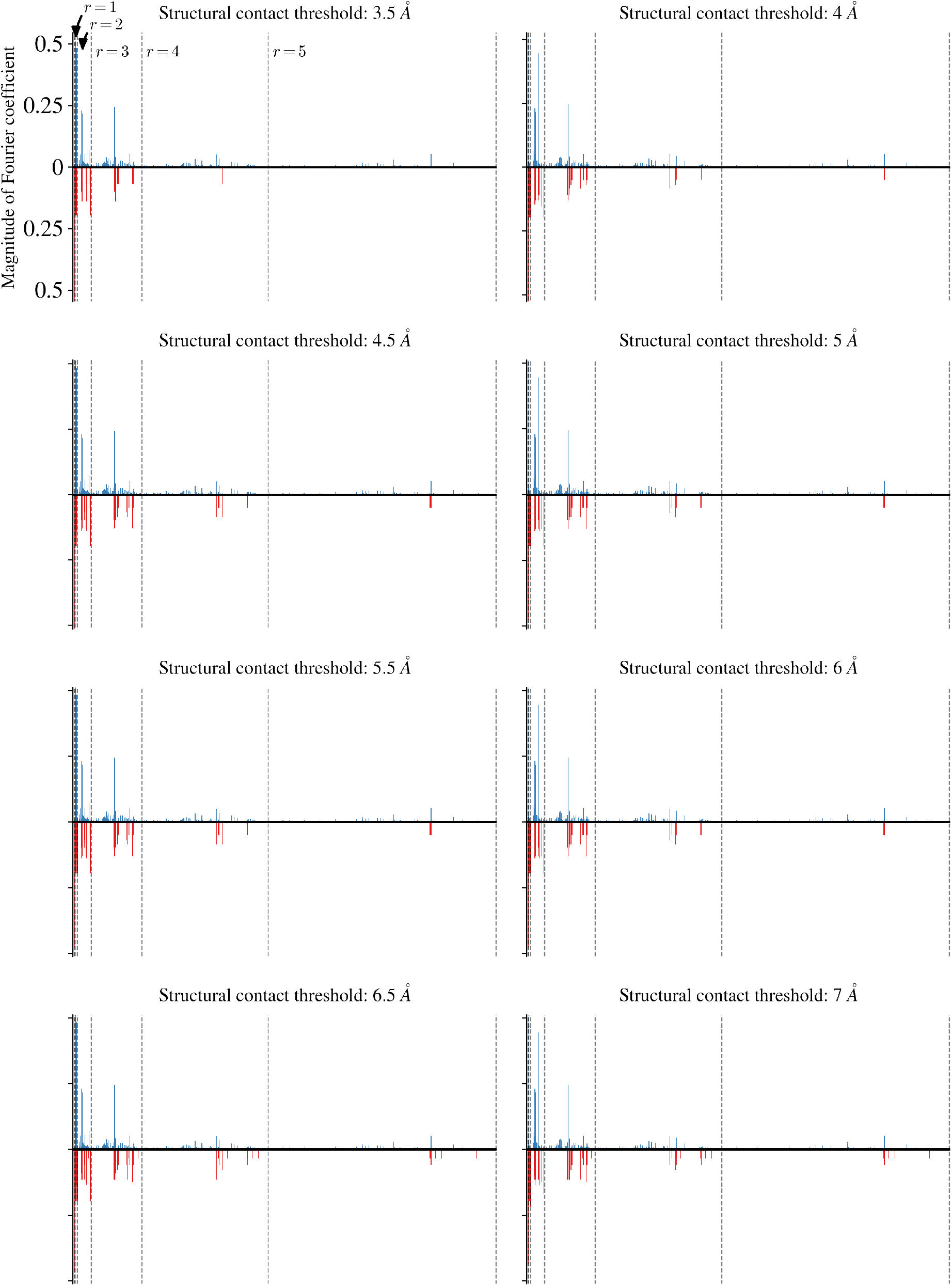
Comparison between magnitudes of Fourier coefficients in the mTagBFP2 empirical fitness function and GNK models with Structural neighborhoods derived from the TagBFP crystal structure at a range of threshold distances. Coloring and details of each plot are as in Fig. 4B of the main text. The title of each plot indicates the threshold distance used to determine Structural neighborhoods of the GNK models whose coefficients are displayed as red bars in the plots.

**Fig. S6:**
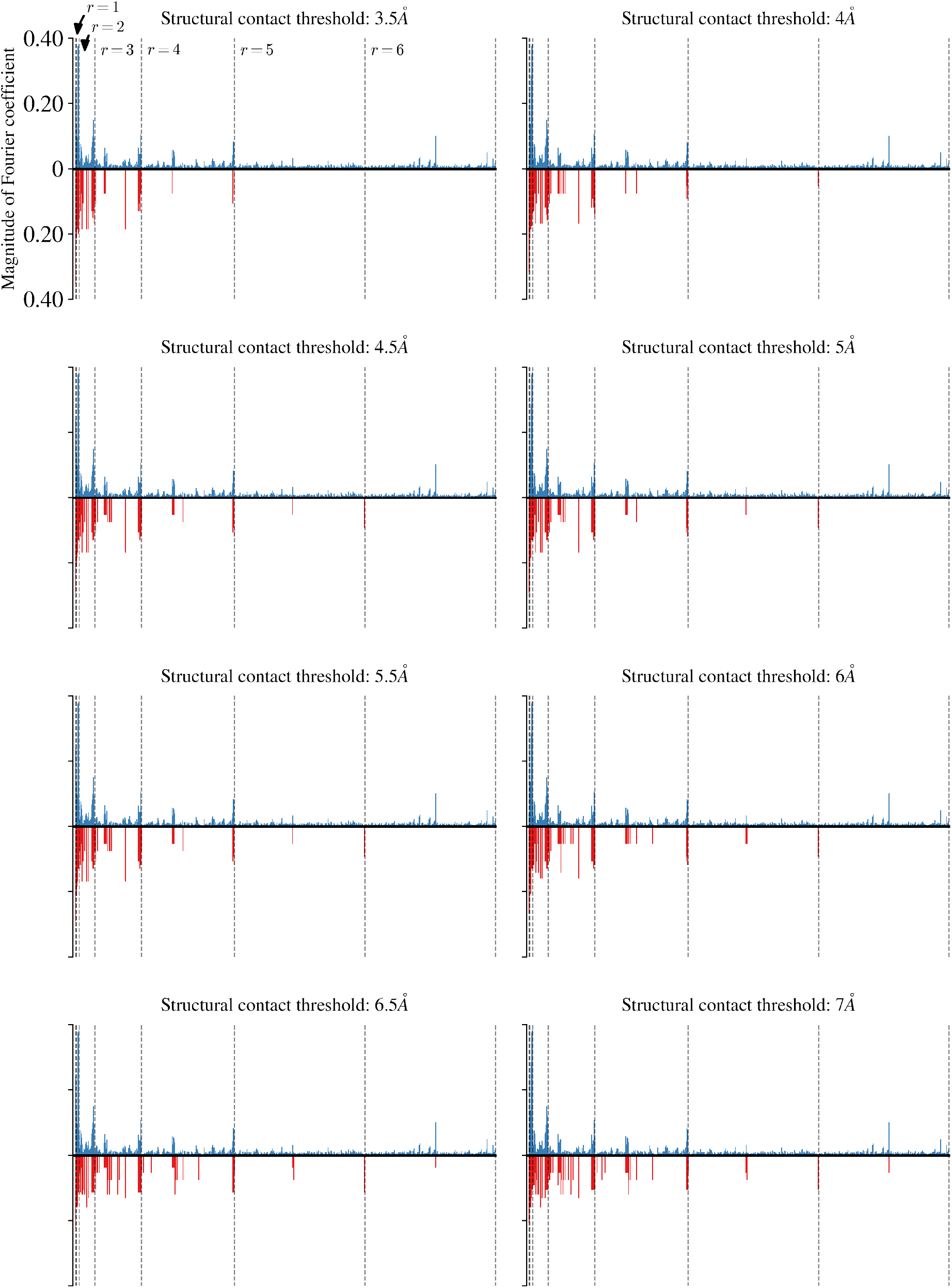
Comparison between magnitudes of Fourier coefficients in the mTagBFP2 empirical fitness function and GNK models with Structural neighborhoods derived from the I-TASSER predicted structure of His3p at a range of threshold distances. Coloring and details of each plot are as in Fig. 4B of the main text. The title of each plot indicates the threshold distance used to determine Structural neighborhoods of the GNK models whose expected coefficient magnitudes are displayed as red bars in the plot.

**Fig. S7:**
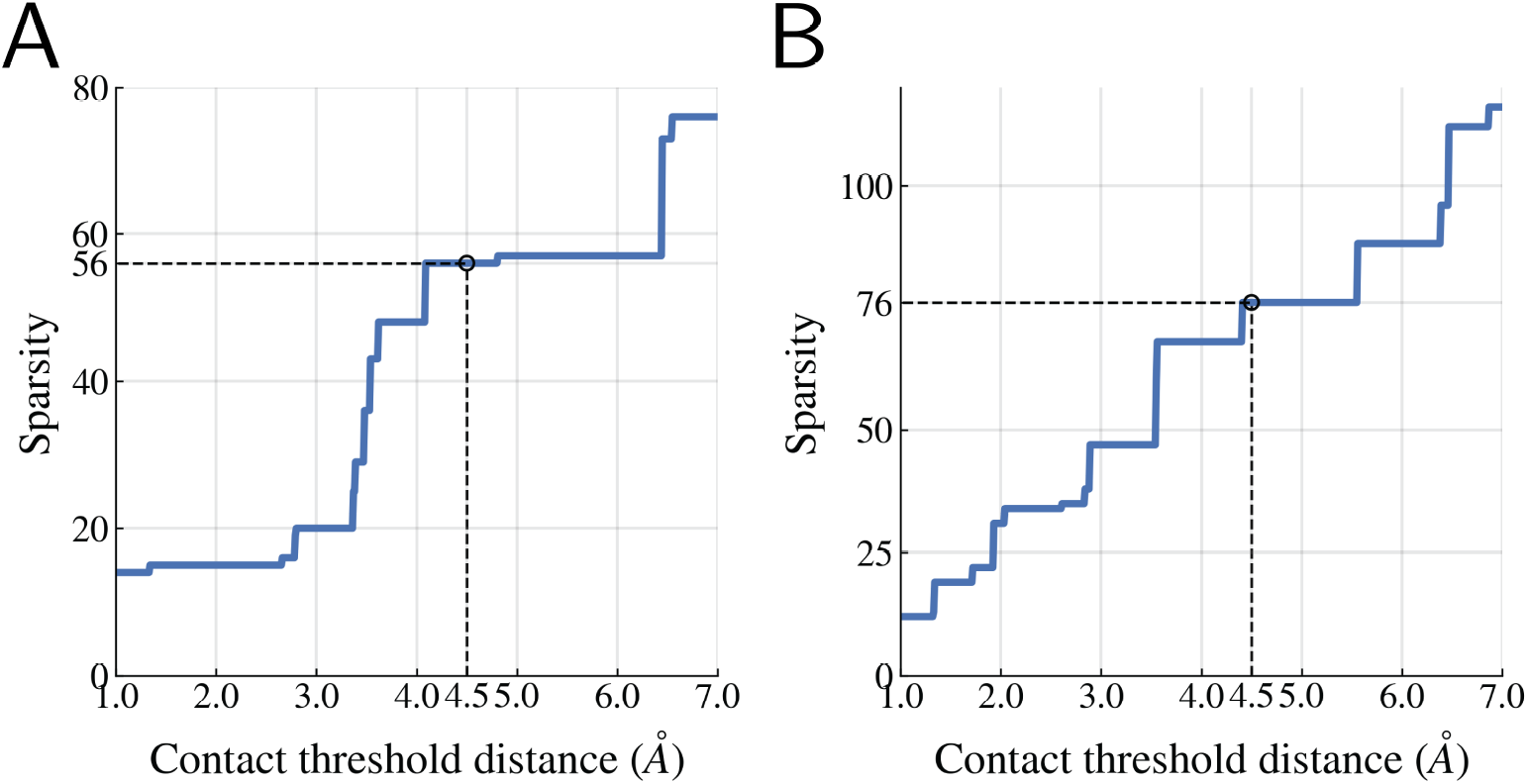
Effect of structural contact threshold distance on sparsity of GNK fitness functions with Structural neighborhoods based on the (A) TagBFP crystal structure and (B) I-TASSER predicted structure of His3p. The horizontal axis of each plot indicates the threshold distance used to define structural contacts and thus used to construct the Structural Neighborhoods. The vertical axis indicates the sparsity of the GNK fitness functions when the Structural Neighborhoods are defined with each threshold distance. The threshold distance used in the main text (4.5Å) and the corresponding sparsities are indicated with dashed lines.

### S5.2 Distribution of largest coefficients by order of interaction

It is not immediately clear from the presentation of Fig. 4B how the largest Fourier coefficients in the empirical and GNK fitness functions are distributed among the orders of epistatic interactions. Fig. S8 shows the fraction of the largest *S* coefficients that correspond to epistatic interactions of each order, with *S* equal to the sparsity of GNK fitness functions.

**Fig. S8:**
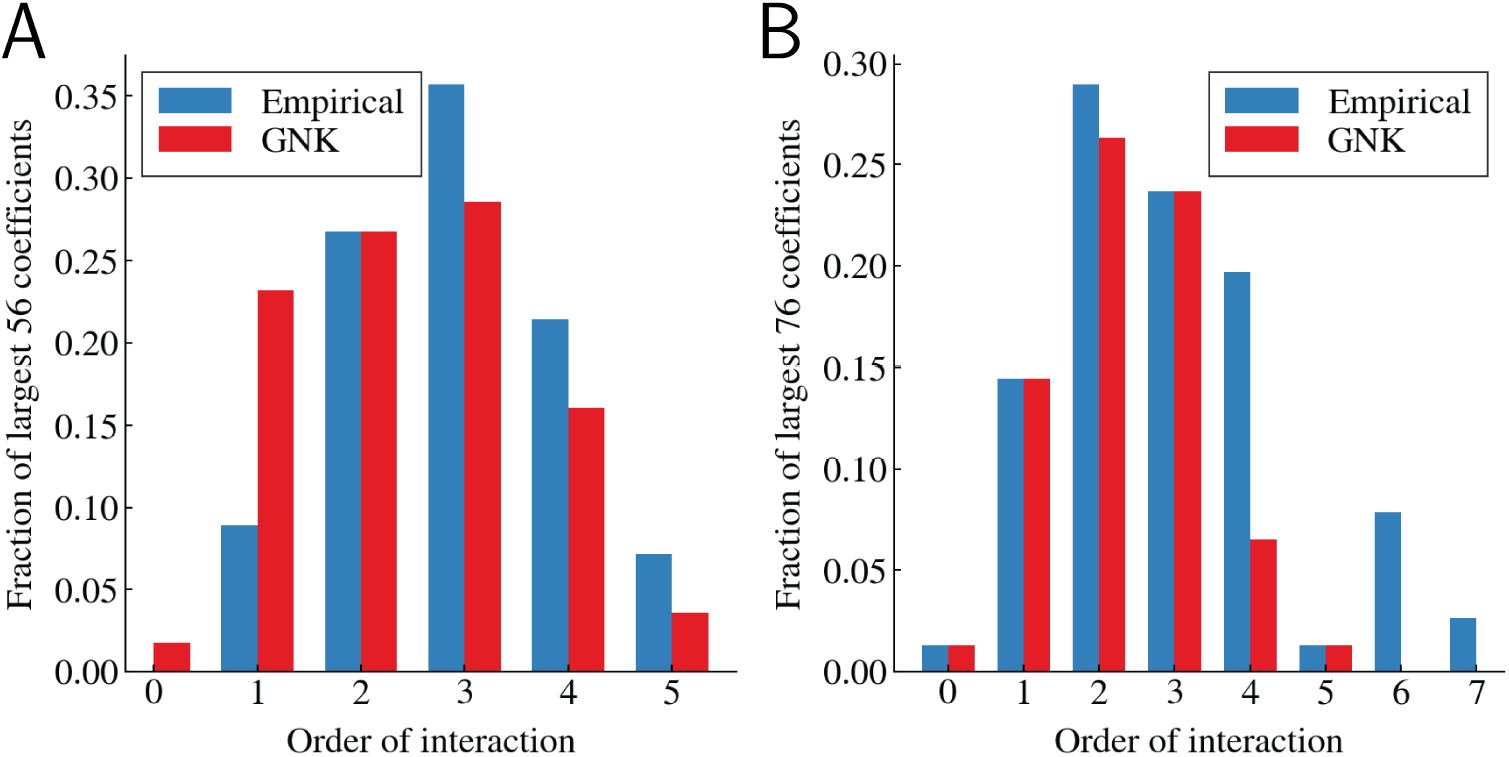
Fraction of Fourier coefficients with the largest magnitudes that are of each order of interactions in the (A) mTagBFP2 empirical fitness function and associated GNK model with Structural neighborhoods and (B) His3p empirical fitness function and associated GNK model with Structural neighborhoods. Blue bars indicate the fraction of the largest *S* empirical coefficients that are of each order of interactions and red bars indicate the fraction of the GNK coefficients with the largest expected magnitudes that are of each order of interactions. *S* is equal to the sparsity of the GNK fitness functions in each case: *S* = 56 in (A) and *S* = 76 in (B).

### S5.3 Analysis of the overlap between empirical and GNK Fourier coefficients

Here we provide more analysis of the overlap between the Fourier coefficients of the empirical protein fitness functions and the corresponding GNK models with Structural neighborhoods than is shown in Fig. 4B.

First, Fig. S9 contains scatter plots that compare the magnitudes of the Fourier coefficients of the empirical protein fitness functions with the expected magnitudes of the corresponding GNK models with Structural neighborhoods. In both cases, we see statistically significant correlation between the expected magnitudes of the GNK coefficients and the magnitudes of the empirical coefficients (*p* = 9.5 × 10^−310^ and *p* = 9.5 × 10^−172^ in the TagBFP and His3p cases, respectively). However, this visualization and quantification technique is not ideal for informing on our claims because it focuses on the coefficients’ magnitude more than the sparsity of the coefficients. Indeed, what is more important for our results is that the GNK model identifies many of the largest empirical coefficients as being nonzero, which is analyzed more concretely in later plots and described below.

In particular, Fig. S10 shows the fraction of the GNK coefficients with the largest expected magnitudes that are also among the empirical coefficients with the largest magnitudes. For example, among the largest 25 coefficients in the TagBFP empirical fitness function, 76% are also among the 25 GNK coefficients with the largest expected magnitudes (which corresponds 19 out of 25 overlapping coefficients). Further, there is 96% overlap between the 25 largest His3p(small) empirical and GNK coefficients (i.e. there are 24 out of 25 overlapping coefficients). At the sparsities of the GNK models (56 and 76 coefficients, respectively, for the TagBFP and His3p(small) cases), there is 41% and 62% overlap between the largest empirical and GNK coefficients in the TagBFP and His3p(small) cases.

Further, we performed statistical tests to determine whether the GNK models with Structural neighborhoods were able to identify the largest coefficients in the empirical fitness functions. To start, we partitioned the empirical coefficients into two sets: those that are identified as being nonzero in the GNK fitness functions and those that are zero in the GNK fitness functions. Kernel Density Estimates (KDEs) of the density of magnitudes of the coefficients in these two sets are shown in Fig. S11. All KDEs were calculated with the Scikit-learn package [62] using a Gaussian kernel with bandwith equal to 0.01. Visually, it appears that the empirical coefficients corresponding to nonzero GNK coefficients indeed tend to be larger than those associated with zero GNK coefficients. To quantify this claim, we performed a Wilcoxon rank-sum test [63] to test the null hypothesis that the two sets of coefficients are sampled from the same population, against the alternative hypothesis that the empirical coefficients corresponding to nonzero GNK coefficients are sampled from a population that is stochastically greater that of the coefficients corresponding to zero GNK coefficients. The p-values resulting from this test are shown in both panels, demonstrating that we can safely reject the null hypothesis in both cases.

We also made these density visualizations and ran the associated statistical test for coefficients corresponding to every order of epistatic interaction where the GNK model has more than one nonzero coefficient (ignoring r=0 and r=1, where the GNK model assigns nonzero variance to all coefficients). Figs. S12 and S13 show these visualizations for the mTagBFP and His3p fitness functions, with the order of interactions indicated by the title of the panels and the p-values of each statistical test displayed in the panels. In all cases, we can reject the null hypothesis at a significance threshold of 0.01.

**Fig. S9:**
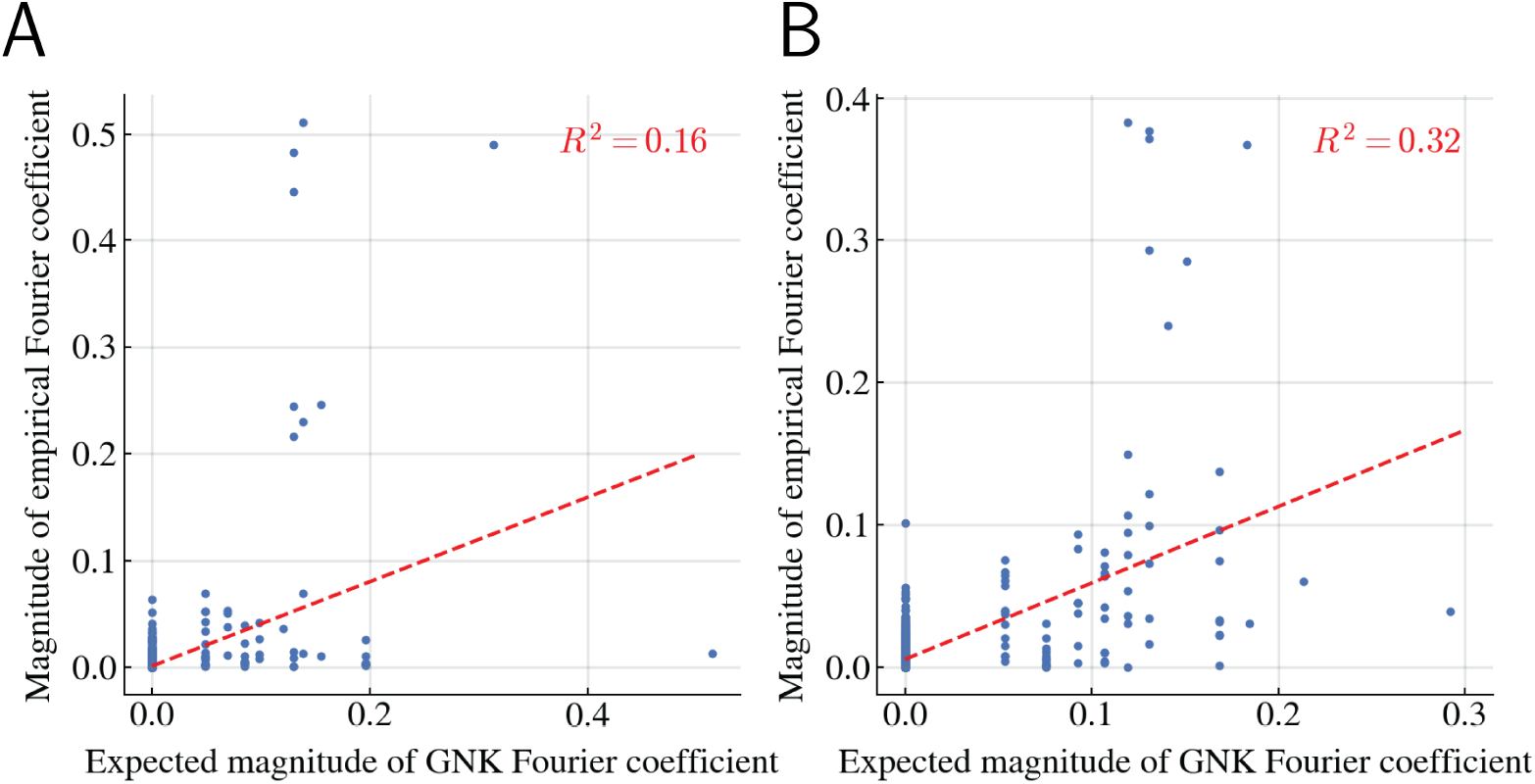
Comparison of the correlation between the magnitudes of the Fourier coefficients of the (A) mTagBFP2 and (B) His3p empirical fitness functions and the expected magnitudes of the Fourier coefficients of the corresponding GNK models with Structural neighborhoods.

**Fig. S10:**
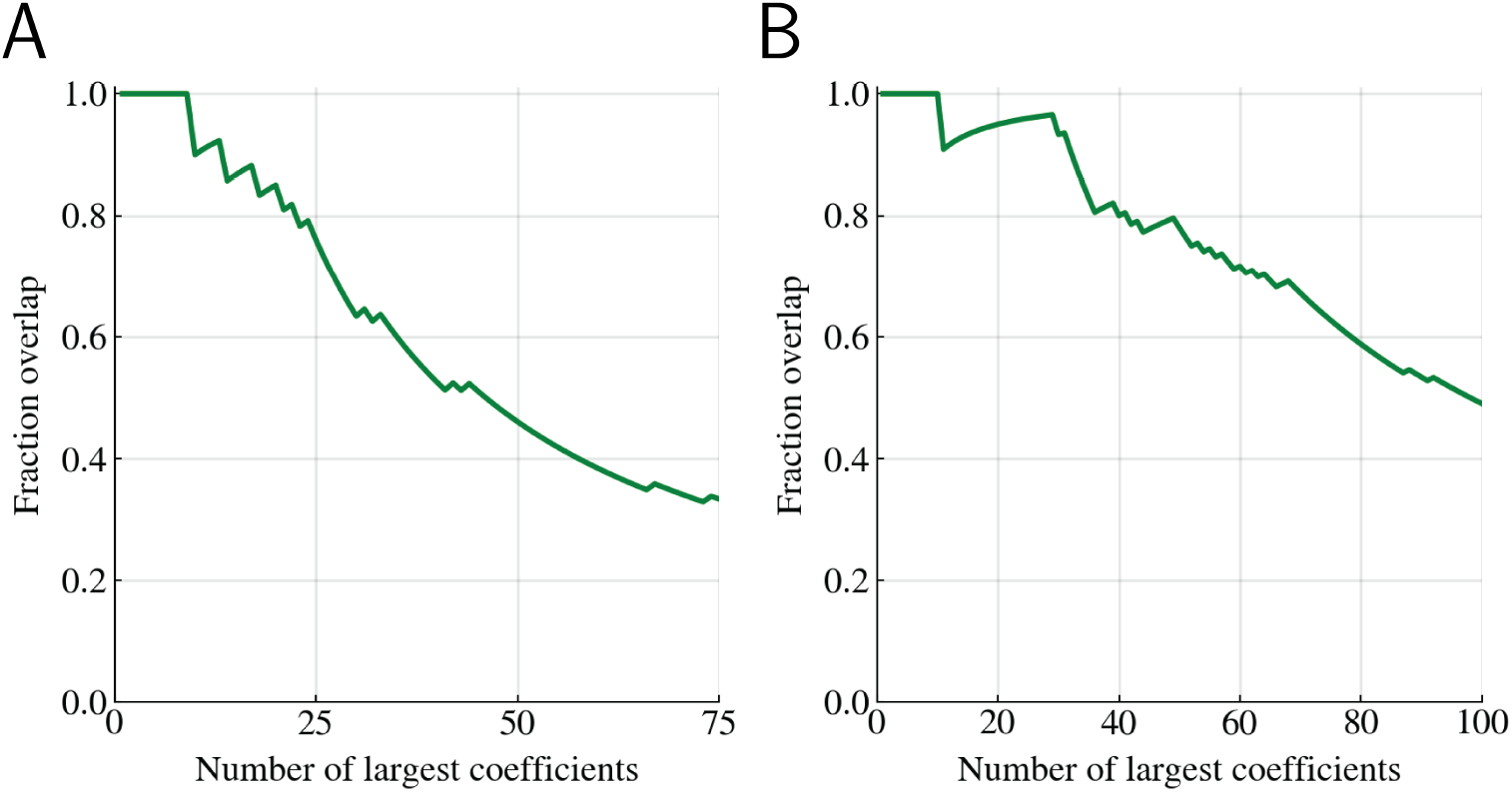
Fraction overlap between Fourier coefficients with largest magnitudes of empirical protein fitness functions and the Fourier coefficients with the largest expected magnitudes in the associated GNK models with Structural neighborhoods. The horizontal axis indicates the number of the largest empirical coefficients that are considered. At a value *S* on the horizontal axis, the vertical indicates the number of the *S* largest empirical coefficients that are also among the *S* coefficients in the GNK model with the largest expected magnitude. The panels correspond to the (A) mTagBFP2 empirical fitness function and associated GNK model with Structural neighborhoods and (B) His3p empirical fitness function and associated GNK model with Structural neighborhoods.

**Fig. S11:**
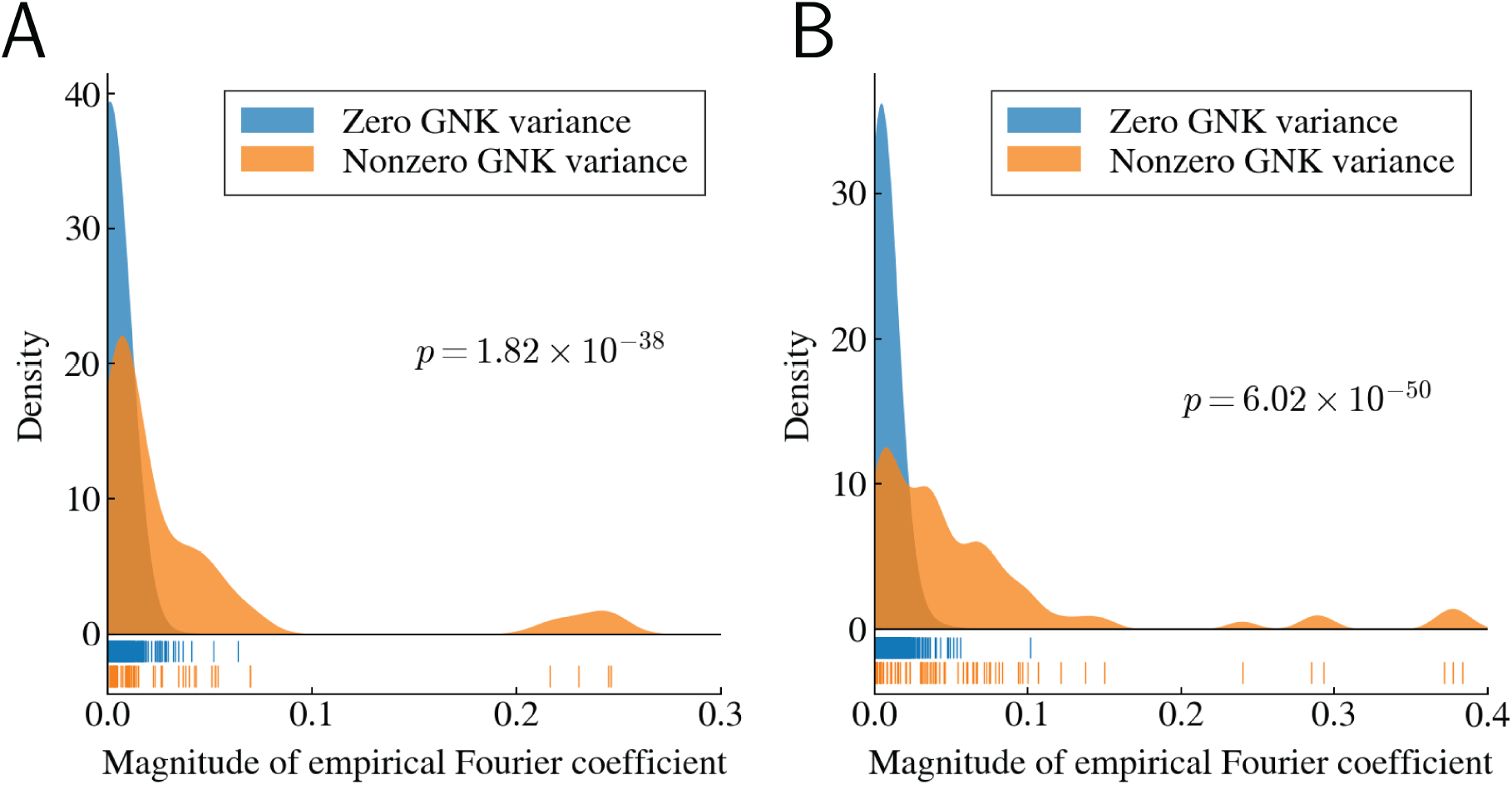
Kernel density estimates of the density of magnitudes of empirical Fourier coefficients that are identified as zero (blue) and nonzero (orange) by GNK models with Structural neighborhoods. The raw magnitudes of the coefficients in each set are shown by the vertical bars below the density plots. The panels correspond to the (A) mTagBFP2 empirical fitness function and (B) His3p empirical fitness function. The p-values associated with a Wilcoxon rank-sum test comparing the two populations of magnitudes in each panel are shown in that panel.

**Fig. S12:**
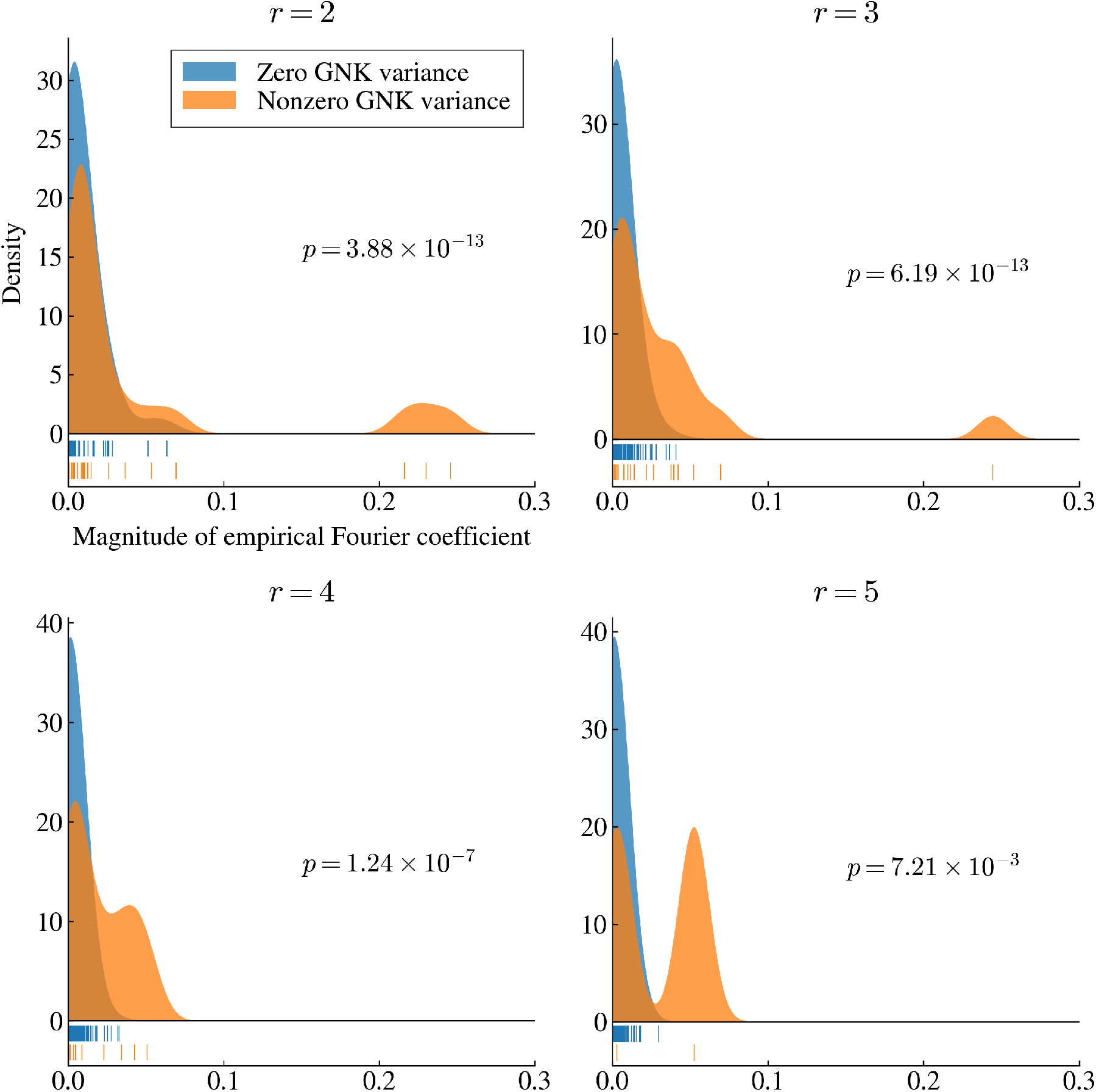
Kernel density estimates of the density of magnitudes of Fourier coefficients in the mTagBFP2 empirical fitness function that are identified as zero (blue) and nonzero (orange) by the associated GNK model with Structural neighborhoods. Each panel corresponds to a particular order of epistatic interaction, indicated by the title of the panel. The p-values associated with a Wilcoxon rank-sum test comparing the two populations of magnitudes in each panel are shown in that panel.

**Fig. S13:**
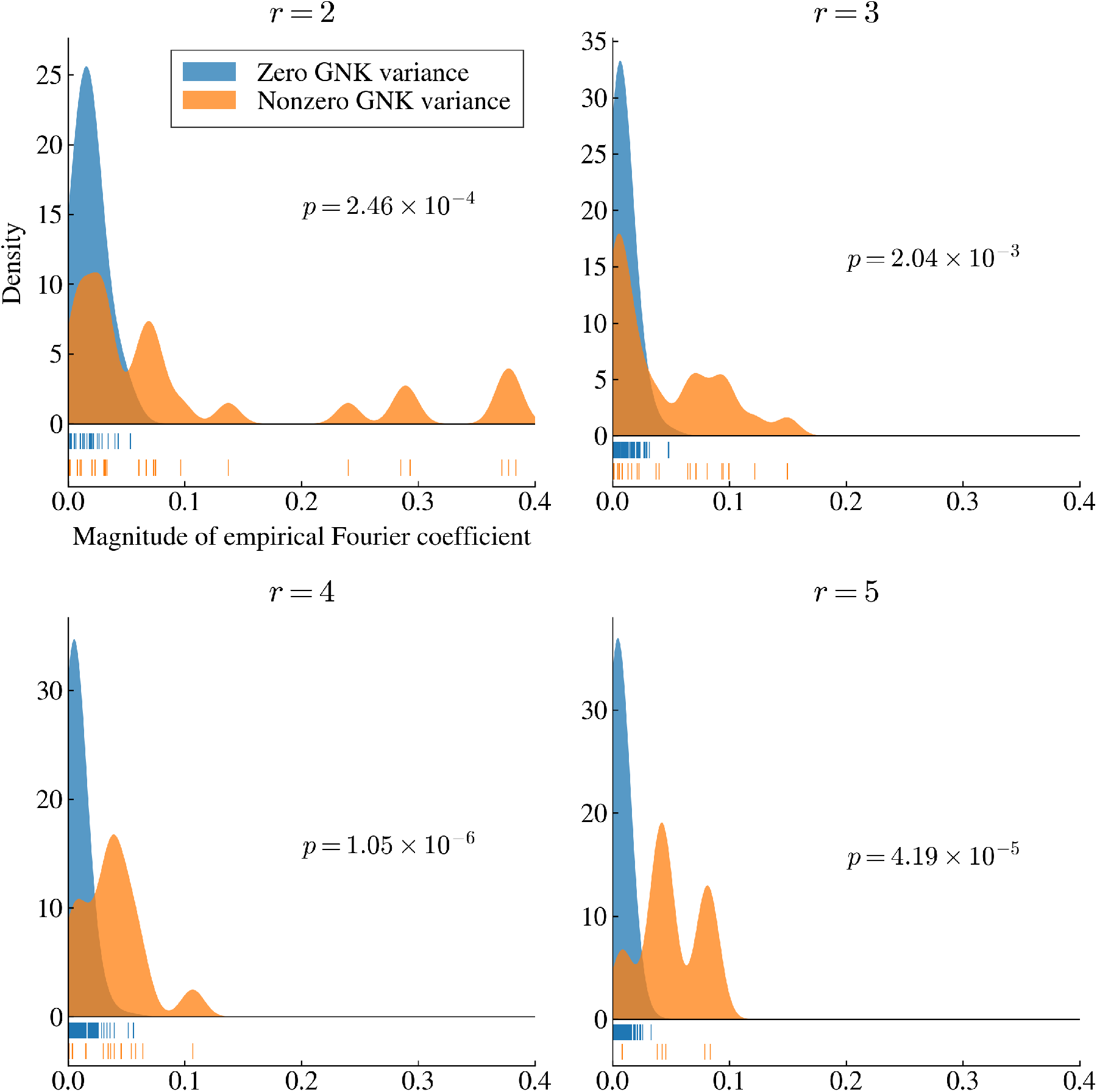
Kernel density estimates of the density of magnitudes of Fourier coefficients in the His3p empirical fitness function that are identified as zero (blue) and nonzero (orange) by the associated GNK model with Structural neighborhoods. Each panel corresponds to a particular order of epistatic interaction, indicated by the title of the panel. The p-values associated with a Wilcoxon rank-sum test comparing the two populations of magnitudes in each panel are shown in that panel.The p-values associated with a Wilcoxon rank-sum test comparing the two populations of magnitudes in each panel are shown in that panel.

### S5.4 Comparison of error bound for GNK and empirical fitness functions

In the main text, we have been primarily concerned with the exact recovery of fitness functions. However, Eq. 8 also provides a means to probe the amount of error that may result from using fewer training samples than are needed for exact recovery. In particular, the function 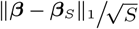 roughly sets the scale for the decay in error as more samples are added (in the sense that it is proportional to the bound on error in the noiseless case). Below we plot this function for the (A) TagBFP and (B) His3p(small) (B) empirical fitness functions, together with the mean and variance of this quantity for 1,000 samples of the corresponding GNK models with Structural neighborhoods (in the same manner that we calculated the percent variance explained curves in Fig. 4C).

**Fig. S14:**
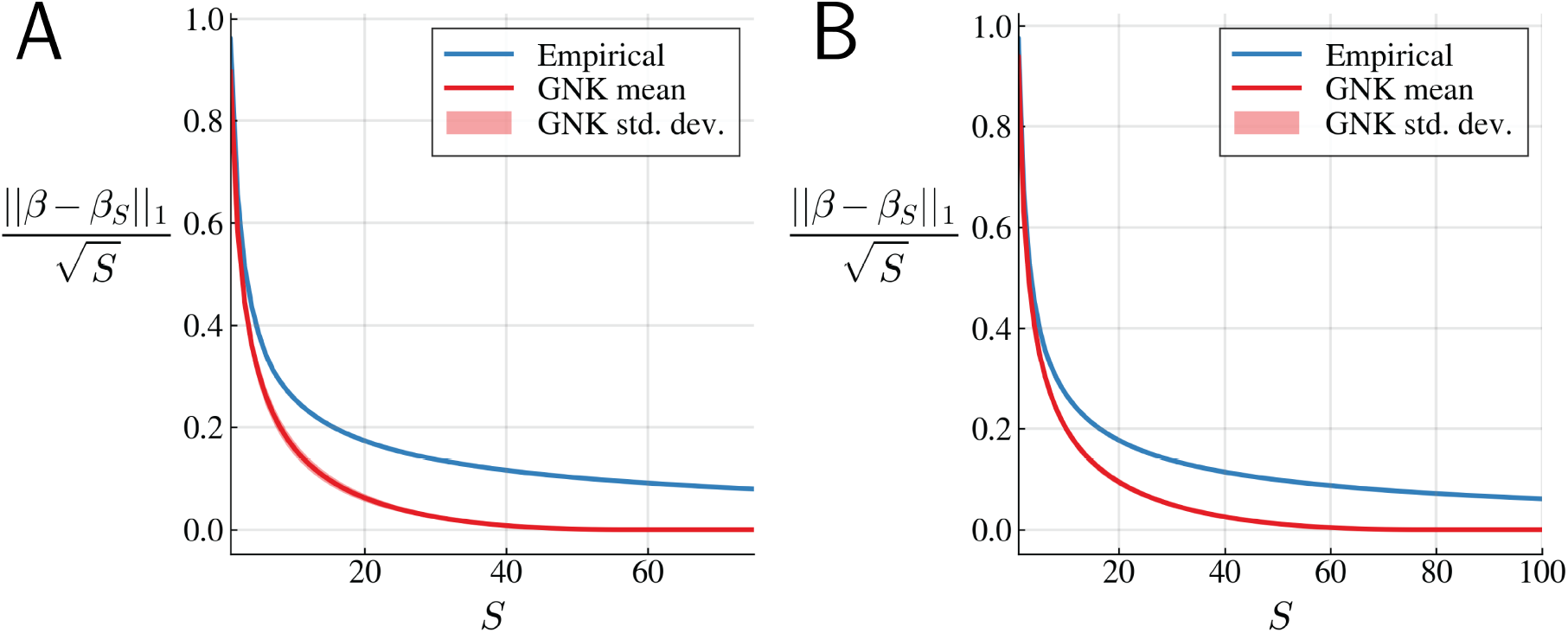
Comparison of the noiseless error bound of Eq. Eq. 8 for empirical protein fitness functions and GNK fitness functions with Structural neighborhoods. Blue curves represent the error bound for the empirical fitness functions, while red curves represent the mean bound of sampled GNK fitness functions, and the red shaded region represent the standard deviation of the bound among these samples. The panels correspond to (A) the mTagBFP2 empirical fitness function and corresponding GNK model and (B) the His3p empirical fitness function and corresponding GNK model. Note that each vector of coefficients has been normalized such that the L1 norm is equal to 1, so that the GNK and empirical coefficients can be compared.

## S6 Proofs and additional theoretical results

In this section, we formally state the mathematical results from the main text and provide proofs.

### S6.1 Graph theory preliminaries

Much of the following requires substantial graph-theoretic construction, so we first introduce the requisite notation and simple definitions. We will use the notation *V*(*G*) and *E*(*G*) to denote the vertex and edge sets of a graph *G*. The graph is then specified by *G* = (*V*(*G*), *E*(*G*)). The “degree” of a vertex *v* is the number of other vertices that are adjacent to *v*. A *k*-regular graph is a graph in which every vertex has degree equal to *k*. The Graph Laplacian of a graph *G* with vertices 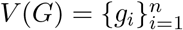 is given by **L**(*G*) ≔ **D**(*G*) – **A**(*G*) where **D**(*G*) is an *n* × *n* diagonal matrix whose *i*^th^ diagonal element is equal to the degree of vertex *i* and **A**(*G*) is the *n* × *n* adjacency matrix of *G* with elements given by

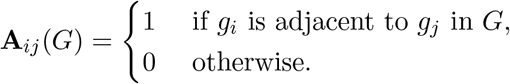

Graph Laplacians and adjacency matrices are real, symmetric matrices and thus have orthonormal sets of eigenvectors. In the case of a *k*-regular graph, **L**(*G*) = *k***I** – **A**(*G*). Thus the Laplacian and adjacency matrices share eigenvectors, and the eigenvalues of the Laplacian are given by λ_*j*_(**L**) = *k* – λ_*j*_(**A**) for *j* = 1, …, *L* where λ_*j*_(**A**) are the eigenvalues of the adjacency matrix.

We will make use of the Cartesian product of graphs, defined below:

#### Definition 1

(Cartesian Product of Graphs). The Cartesian product between two graphs *G* = (*V*(*G*), *E*(*G*)) and *H* = (*V*(*H*), *E*(*H*)) is defined as *G*□*H* = (*V*(*G*) × *V*(*H*), *E*(*G*□*H*)), where two vertices (*g*, *h*) and (*g*′, *h*′) are adjacent in *G*□*H* if and only if either

1. *g* = *g*′ and *h* is adjacent to *h*′ in *H*, or
2. *h* = *h*′ and *g* is adjacent to *g*′ in *G*.

A direct consequence of Definition 1 is that the adjacency matrix of the Cartesian product can be constructed from the adjacency matrices of its components as [64]:

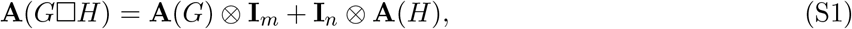

where *m* = |*V*(*H*)| and *n* = |*V*(*G*)| are the number of vertices in *H* and *G*, respectively.

We will additionally make use of the Lexicographic product of graphs [57].

#### Definition 2

(Lexicographic Product of Graphs). The Lexicographic product between two graphs *G* = (*V*(*G*), *E*(*G*)) and *H* = (*V*(*H*), *E*(*H*)) is defined as *G* ∘ *H* = (*V*(*G*) × *V*(*H*), *E*(*G* ∘ *H*)), where two vertices (*g, h*) and (*g*′, *h*′) are adjacent in *G* ∘ *H* if and only if either

1. *g* is adjacent to *g*′ in *G*, or
2. *g* = *g*′ and *h* is adjacent to *h*′ in *H*.

The adjacency matrix of a Lexicographic product of graphs is given by [64]:

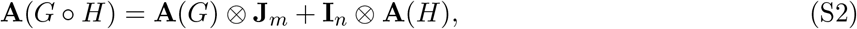

where **J**_*m*_ is the *m* × *m* matrix with every element equal to one.

The graphs described up until now have been ‘simple’ graphs, where each edge connects exactly two vertices and vertices are connected by at most one edge. We will also discuss ‘hypergraphs’, where ‘edges’ are sets that may contain more than two vertices (these are referred to as ‘hyperedges’). Let *H* = (*V*(*H*), *E*(*H*)) be a hypergraph with *n* vertices, 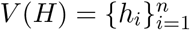, and *p* hyperedges, 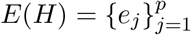. The *incidence* matrix of *H*, denoted **F**(*H*),9 is the *n* × *p* matrix with elements

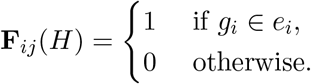

The degree of a vertex in a hypergraph is equal to the number of hyperedges that contain that vertex, and a *k*-regular hypergraph is one in which all vertices have degree equal to *k*. The *clique multigraph* corresponding to a hypergraph is the multigraph (another extension of simple graphs where two vertices can have multiple simple edges between them) with the same vertices as the hypergraph, and as many edges between two vertices as the number of times those vertices co-occur in a hyperedge of the hypergraph (i. e., if two vertices are both in two separate hyperedges of the hypergraph, then they will have two edges between them in the clique multigraph) [65]. The (*i, j*)^th^ element of the adjacency matrix of a multigraph is equal to the number of edges that connect the *i*^th^ and *j*^th^ vertices.

Below are two Lemmas regarding the spectrum of Cartesian and lexicographic graph products that will be useful [66].

#### Lemma 1.

*Let G and H be regular graphs with n and m vertices, respectively. Let* **A**(*G*) = **PΛ***_G_***P**^*T*^ *and* **A**(*H*) = **QΛ**_*H*_**Q**^*T*^ *be eigendecompositions of the adjacency matrices of G and H, respectively. Then the adjacency matrix of the Cartesian product G*□*H has the eigendecomposition given by*:

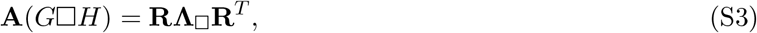

*where* **R** = **P** ⊗ **Q** *and* **Λ**_□_ = **Λ**_*G*_ ⊗ **I***_m_* + **I**_*n*_ ⊗ **Λ***_H_*.

*Proof*. We can use Equation S1 to prove the proposed Lemma directly:

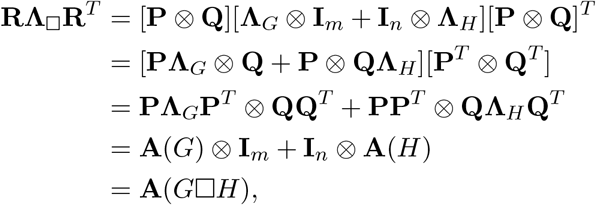

where in the second and third lines we have used the property of Kronecker products that (**A** ⊗ **B**)(**C** ⊗ **D**) = (**AB** ⊗ **CD**), in the second line we have also used the fact that the transpose is distributive over the Kronecker product, (**C** ⊗ **D**)^*T*^ = **C**^*T*^ ⊗ **D**^*T*^, and the final line is a result of Equation S1.

#### Lemma 2.

*Let G and H be regular graphs with n and m vertices, respectively. Let* **A**(*G*) = **PΛ**_*G*_**P**^*T*^ *and* **A**(*H*) = **QΛ**_*H*_**Q**^*T*^ *be eigendecompositions of the adjacency matrices of G and H*. *Then the adjacency matrix of the lexicographic product G* ∘ *H has the eigendecomposition given by*:

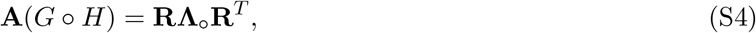

*where* **R** = **P** ⊗ **Q**, **Λ**_∘_ = **Λ**_*G*_ ⊗ **B** + **I**_*n*_ ⊗ **Λ**_*H*_, *and* 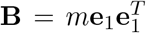 (*i. e*., *B_ij_* = *m if i* = *j* = 1 *and zero otherwise*).

*Proof*. The adjacency matrix of any *k*-regular graph with *m* vertices has two eigenvalues, *k* and 0, with multiplicities 1 and *m* – 1, respectively. The normalized eigenvector associated with the eigenvalue *k* is 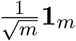 and the normalized eigenvectors associated with the eigenvalue 0 are any set of length-*m* orthogonormal vectors that are orthogonal to **1**_*m*_ (i. e., vectors that sum to one). Since *H* is a regular graph, we then have that

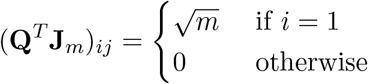

and

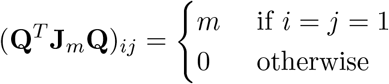

Therefore, (**Q**^*T*^ **J**_*m*_**Q**) = **B** and further **QBQ**^*T*^ = **J**_*m*_. Then we can use Eq. S2 to show:

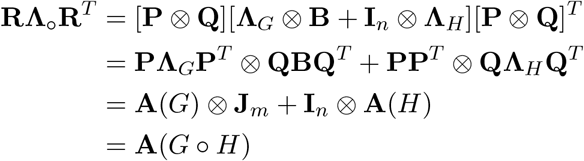

### S6.2 Graph Fourier basis results

Here we will prove results related to our construction of Graph Fourier bases. First, we prove a result regarding the eigenvectors of the complete graph, which is presented in the main text as Eq. 9.

#### Proposition 1.

*An orthonormal set of eigenvectors of the Graph Laplacian of the complete graph K*(*q*) *are given by the columns of the q* × *q matrix*:

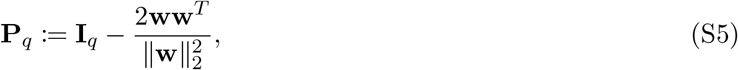

*where* 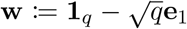, **1**_*q*_ *is the vector of length q whose elements are all equal to one*, **e**_1_ *is the length q with the first element set to 1 and all others set to zero, and* **I**_*q*_ *is the q* × *q identity matrix*

*Proof of Proposition 1*. The Graph Laplacian of the complete graph *K*(*q*) is given by **L**(*K*(*q*)) = *q***I**_*q*_ – **J**_*q*_ (i. e., the *q* × *q* matrix with *q* on the diagonal and all other elements equal to –1). This matrix has two eigenvalues, 0 and *q*, with multiplicities 1 and *q* – 1, respectively. The normalized eigenvector corresponding to the zero eigenvalue is 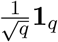. Since the graph Laplacian is symmetric, the eigenvectors are orthogonal and therefore the remaining eigenvectors are any set of *n* – 1 orthogonal vectors that are orthogonal to **1**_*q*_ (i.e., vectors that sum to zero) and each other. In order to show that the columns of the Householder matrix given in Eq. S5 are orthonormal eigenvectors of the complete graph, we will prove (i) that the first column of **P**_*q*_ is equal to 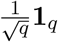 and (ii) that **P**_*q*_ is an orthogonal matrix:

i. We can more explicitly write the **w** = [*w*_1_, *w*_2_, …, *w*_*q*_] vector as 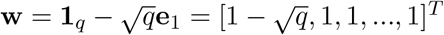. Therefore, 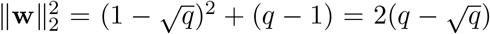. Let *α_i_* for *i* = 1, 2, .. *q* be the elements of the first column of **P**_*q*_, respectively. Then,

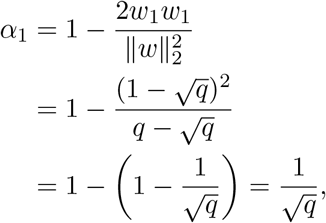

and for *j* = 2, 3, …, *q*, we have

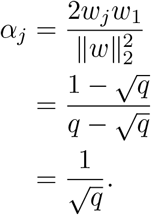 Therefore, all elements of the first columns of **P**_*q*_ are equal to 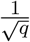.
ii. The orthogonality of **P***q* follows directly from the definition:

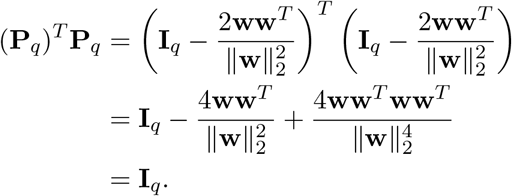

Thus, **P**_*q*_ is an orthogonal matrix whose first column is 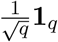, and further, the columns are **P**_*q*_ are an orthonormal set of eigenvectors of *K*(*q*).

It is worth noting that the adjacency matrix of the complete graph is an example of a circulant matrix, and thus an alternative basis to that of Eq. S5 is the *q*-point Discrete Fourier Transform [67]. Using the DFT matrix along with Eq. 10 results in the basis for the Hamming graph presented in [68].

We additionally have the following result showing how to construct the Graph Fourier basis corresponding to the Hamming graph using the eigenvectors of the complete graph, which is presented in the main text as Eq. 10.

#### Proposition 2.

*An orthonormal set of eigenvectors of the Graph Laplacian of the Hamming graph H*(*L*, *q*) *are given by the columns of the q^L^ × q^L^ matrix*

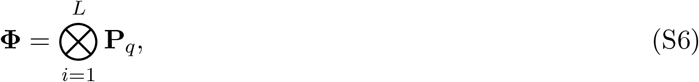

*where* **P**_*q*_ *is defined in* Eq. S5.

In order to prove Proposition 2, we need a preliminary result. First remember that the Hamming graph *H*(*L*, *q*) is the *L*-fold Cartesian product of the complete graph *K*(*q*). We have the following result regarding the eigenvectors and eigenvalues of the adjacency matrices of Cartesian products of regular graphs.

#### Lemma 3.

*Let G and H be regular graphs and let* **P** *and* **Q** *be matrices whose columns are eigenvectors of the Graph Laplacians of G and H*, *respectively. Then the columns of* **P** ⊗ **Q** *are eigenvectors of the Graph Laplacian of the Cartesian product G*□*H*.

*Proof*. For a regular graph, the degree matrix is a constant multiplied by the identity matrix. Thus, in this case, the Graph Laplacian and adjacency matrices differ only by a constant added to the diagonal (and a constant multiplicative factor of –1). The Graph Laplacian and adjacency matrices of a regular graph therefore have the same eigenvectors. The result then follows from Lemma 1.

*Proof of Proposition 2*. The Hamming graph *H*(*L,q*) is defined as the *L*-fold Cartesian product of the complete graph *K*(*q*) [56]:

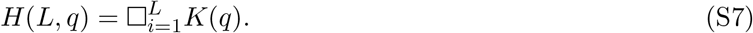

Thus, by Lemma 3, the eigenvectors of the Graph Laplacian of *H*(*L,q*) are the *L*-fold Kronecker product of the eigenvectors of the Graph Laplacian of *K*(*q*), as given in Eq. S6.

### S6.3 Distribution of GNK Fourier coefficients

Here we prove our result regarding the distribution of the Fourier coefficients of fitness functions sampled from the GNK model (Eq. 5). To begin, let 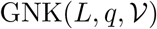 be probability distribution over fitness functions induced by the GNK model for sequence length *L*, alphabet size *q*, and a set of neighborhoods corresponding to each position, 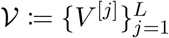. We now formally restate the result of Eq. 5.

#### Theorem 1.

*Let* 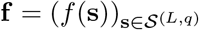 *be the complete vector of evaluations of a fitness function* 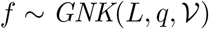. *Then the Fourier coefficients of f, given by **β*** = **Φ**^*T*^**f**, *are distributed according to* 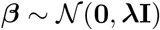 (*i. e., normally distributed with zero mean and diagonal covariance). Let **β**_U_ be the length* (*q* – 1)^*r*^ *sub-vector of **β** representing the epistatic interaction *U*. Then the variance of every element of **β**_U_ is given by*

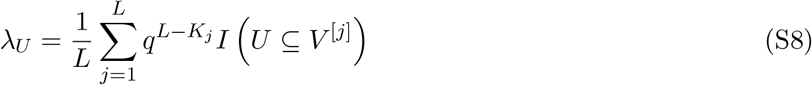

*where I*(*U* ⊆ *V*^[*j*]^) *is an indicator function that is equal to one if U is a subset of or equal to V*^[*j*]^ *and zero otherwise*.

The proof of Theorem 1 is quite involved and requires a number of lemmas; the proofs of the lemmas are shown after the proof of the main result.

In order prove Theorem 1, we will first provide an alternative definition of the GNK model in terms of hypergraphs. To start, we’ll now assign an index to every sequence in the space of sequences, so 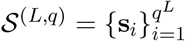. As in the main text, 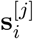 refers to the subsequence of **s**_*i*_ corresponding to the indices in the neighborhood *V*^[*j*]^. Each neighborhood in the GNK model induces a hypergraph over sequence space, where the vertices represent all sequences in 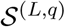 and edges contain sequences that share subsequences corresponding to the indices in the neighborhood. We formally define this hypergraph and related quantities below.

#### Definition 3

(GNK hypergraph). *Let* 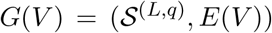 *be a ‘GNK hypergraph’ corresponding to a neighborhood V for a GNK model defined for sequences of length L and alphabet size q. The edge set, E*(*V*), *corresponds to every possible subsequence of length* |*V*|, *and two sequences co-occur in an edge if and only if they share the subsequence corresponding to the positions in V*. *Additionally, let* **F**(*V*) ≔ **F**(*G*(*V*)) *be the incidence matrix of G*(*V*), *C*(*V*) *be the clique multigraph of G*(*V*) *and* **A**(*V*) ≔ **A**(*C*(*V*)) *be the adjacency matrix of C*(*V*). *Finally, when it is appropriate to consider the indexed neighborhoods V*^[*j*]^, *then we will use this indexing for all of the GNK hypergraph quantities. Specifically, define G*^[*j*]^ ≔ *G*(*V*^[*j*]^), **F**^[*j*]^ ≔ **F**(*V*^[*j*]^), *C*^[*j*]^ ≔ *C*(*V*^[*j*]^), *and* **A** ^[*j*]^ ≔ **A**(*V*^[*j*]^).

The following Lemma gives an immediate useful result of this definition.

#### Lemma 4.

*Every GNK hypergraph, G*(*V*), *is a 1-regular hypergraph*.

We will use the GNK hypergraphs to provide an alternative definition of the GNK model, which is shown in the following result. Note that this definition is equivalent to the matrix definition of the GNK model of ref. 30.

#### Lemma 5.

*Define the matrix* **F** *as the column-wise concatenation of the incidence matrices* **F**^[*j*]^ *for j* = 1, 2, …, *L*:

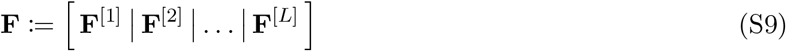

*Additionally, let* 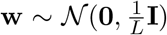 *be a length* 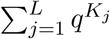 *normally distributed random vector. Then*, **f** = **Fw** *contains all fitness evaluations of a fitness function f that is distributed according to* 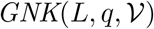 (*i. e*., 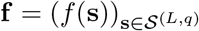 *and* 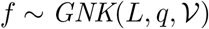).

The following conclusions regarding the statistics of the Fourier coefficients in the GNK model are immediate from this definition of the model. In what follows, we use angled brackets, 〈〉 to indicate expectation over the random field.

#### Lemma 6.

*Let **β** be the Fourier coefficients of a fitness functions distributed according to* 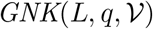. *Then **β** is normally distributed with* 〈***β***〉 = 0 *and covariance matrix*

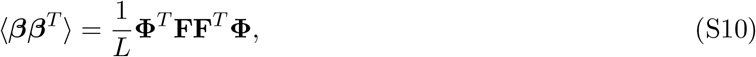

*where* **Φ** *is the Fourier basis defined in* Eq. S6 *and* **F** *is the column-wise concatenation of the incidence matrices of GNK hypergraphs defined in* Eq. S9.

Lemma 6 provides a straightforward path towards proving Theorem 1. We now need to show that (i) **Φ** diagonalizes **FF**^*T*^ (i.e., **Φ** is a basis of eigenvectors for **FF**^*T*^) and (ii) that the eigenvalues of **FF**^*T*^ are given by Eq. S8. The problem can be further simplified by first noting the following simple result, which follows straightforwardly from the multiplication of block matrices.

#### Lemma 7.

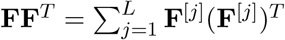.

This result tells us that if possible, it is sufficient to prove that **Φ** diagonalizes each **F**^[*j*]^ (**F**^[*j*]^)^*T*^ in order to prove that **Φ** diagonalizes **FF**^*T*^. Then the eigenvalues of **FF**^*T*^ will simply be given by the sum of the eigenvalues of the **F**^[*j*]^ (**F**^[*j*]^)^*T*^. We are further assisted by the following result regarding the outer product of incidence matrices of regular hypergraphs, due to [65].

#### Lemma 8.

*Let C be the clique multigraph of a *k*-regular hypergraph H with incidence matrix* **F**(*H*). *Then* **F**(*H*)**F**(*H*)^*T*^ = **A**(*C*) + *k***I**, *where* **A**(*C*) *is the adjacency matrix of C*.

Lemma 8 tells us if we can determine the spectrum of the adjacency matrices **A**^[*j*]^ of the clique multigraphs *C*^[*j*]^, then it is straightforward to calculate the spectrum of **F**^[*j*]^ (**F**^[*j*]^)^*T*^. In order to begin to calculate the spectrum of **A**^[*j*]^ we recognize the following simple fact regarding these clique multigraphs (remember that *G*^[*j*]^ is a 1-regular hypergraph by Lemma 4).

#### Lemma 9.

*The clique multigraph of a 1-regular hypergraph is a simple graph*.

We thus need to determine the spectrum of the simple graphs, *C*^[*j*]^. *C*^[*j*]^ contains edges between any two sequences that share a subsequence corresponding to the indices in the *j*^th^ neighborhood *V*^[*j*]^. In order to calculate the spectrum of *C*^[*j*]^, we will first show how these clique multigraphs can be constructed recursively. In the next few Lemmas, we will provide results for clique graphs associated with a generic neighborhood *V*, and then return to considering the indexed neighborhoods *V*^[*j*]^ when necessary.

#### Lemma 10.

*Let V* ⊆ {1, 2, …, *L*} *be a GNK neighborhood. Additionally, let O*(*q*) *be the empty graph of size q (i. e., the graph containing q vertices and no edges) and define the graphs B_l_*(*V*), *via the recursion relation*:

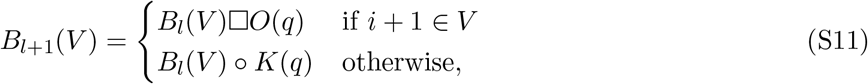

*for i* = 1, 2, …, *L* – 1, *where*

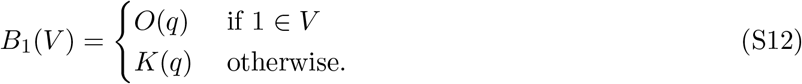

*Then the vertices of B_l_*(*V*) *represent all sequences in* 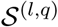 *and two sequences* 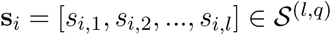 *and* 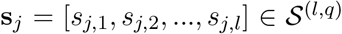 *are adjacent in B_l_*(*V*) *if and only if s_i,k_* = *s_j,k_ for every k* ∈ *V*_(*l*)_ *where we define V*_(*l*)_ ≔ {*m* ∈ *V*: *m* ≤ *l*} *to be the *l* smallest elements of V*.

The simple corollary of Lemma 10 is that the clique graphs *C*(*V*) are the final results of the recursion in Eq. S11.

#### Lemma 11.

*Let C*(*V*) *be the clique multigraph of a GNK hypergraph G*(*V*) *and B_L_*(*V*) *be the graph defined by Equations* Eq. S11 *and* Eq. S12. *Then C*(*V*) = *B_L_*(*V*).

We have thus given a recursive definition of *C*(*V*), which will allow us to calculate the spectrum of the adjacency matrix **A**(*V*) using the spectral properties of graph products presented in Lemmas 1 and 2. In particular, we have the following result regarding the eigenvectors of **A**(*V*).

#### Lemma 12.

*The columns of the Fourier basis* **Φ** *are a complete set of orthonormal eigenvectors of the adjacency matrix* **A**(*V*) ≔ **A**(*C*(*V*)) *of the clique multigraph C*(*V*).

We could similarly use Lemmas 1 and 2 to calculate the eigenvalues of **A**(*V*); however, this would not allow us to connect the eigenvalues to epistatic interactions, as is required to prove Theorem 1. We will instead proceed by showing in Lemma 13 that the columns of suitably defined matrix are eigenvectors of the adjacency matrix **A**(*V*) with eigenvalues equal to a summand of Eq. S8 up to additive constant. Then, in Lemma 14, we will show that this matrix is indeed equal to the columns of the Fourier basis corresponding to the epistatic interaction *U*. For the following results, recall from the main text that 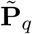 is the matrix containing the final *q* – 1 unnormalized columns of **P**_*q*_, such that 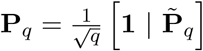, where | denotes column-wise concatenation.

#### Lemma 13.

*Let U* ⊆ {1, 2, …, *L*} *be a set of position indices representing an epistatic interaction and V* ⊆ {1, 2, …, *L*} *be a GNK neighborhood. Define the matrix* **Z**_*l*_(*U*) *with the recursion relation*:

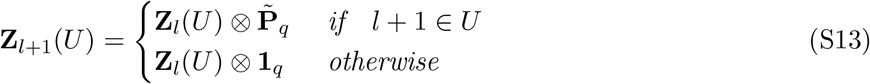

*for l* = 1, 2, …, *L* – 1, *where*

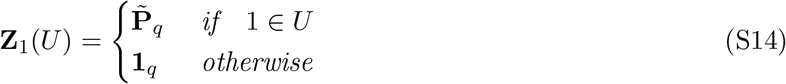

*Then the columns of* **Z**_*L*_(*U*) *are eigenvectors of the adjacency matrix* **A**(*V*), *all associated with the eigen-value given by*

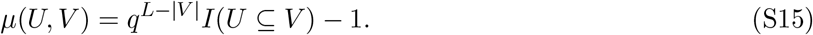

#### Lemma 14.

*Define* **Φ**_*U*_ *as the matrix of* (*q* – 1)^|*U*|^ *columns of the Fourier basis* **Φ** *corresponding to the epistatic interaction U* ⊆ {1, 2, …, *L*}:

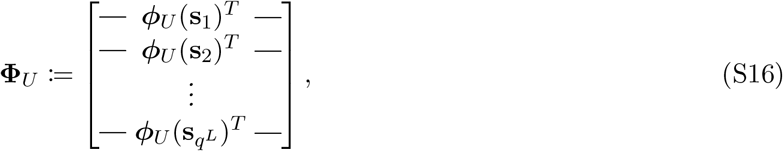

*where* 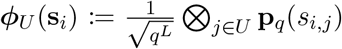 *is the encoding of sequence* **s**_*i*_ *in terms of the epistatic interaction U in the Fourier basis. Then*, 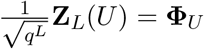, *where* **Z**_*L*_(*U*) *is defined by Equations* Eq. S13 *and* Eq. S14.

Equipped with these results, we are finally prepared to prove Theorem 1.

*Proof of Theorem 1*. In order to prove this theorem, we need to show (i) that the Fourier coefficients are normally distributed with zero mean and diagonal covariance and (ii) that the variance of the coefficients corresponding to a particular epistatic interaction are given by Eq. 5.

First, Lemma 6 proves that the Fourier coefficients are normally distributed with zero mean. Next, Lemma 12 proves that the Fourier basis **Φ** diagonalizes the adjacency matrix **A**^[*j*]^ of the clique multigraph of the GNK hypergraph *G*^[*j*]^. Recalling that *G*^[*j*]^ is a 1-regular hypergraph, then by Lemma 8, **F**^[*j*]^(**F**^[*j*]^)^*T*^ = **A**^[*j*]^ + **I**. Therefore, the Fourier basis diagonalizes **F**^[*j*]^(**F**^[*j*]^)^*T*^ for all *j* = 1, 2, …, *L*, and thus also diagonalizes **FF**^*T*^ due to Lemma 7. Then, the covariance matrix of the Fourier coefficients, which is shown in Lemma 6 to be 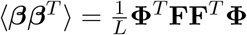, is diagonal. The eigenvalues of **FF**^*T*^ are then equal to the variances of the Fourier coefficients.

Lemma 13 shows that the columns of the matrix **Z**_*L*_(*U*) defined by Equations Eq. S13 and Eq. S14 are eigenvectors of **A**^[*j*]^. Further, Lemma 14 shows that this matrix is equal to the columns of the Fourier basis corresponding to the epistatic interaction *U*, **Φ**_*U*_. Thus, Lemma 13 shows that the eigenvalue of **A**^[*j*]^ associated with the columns **Φ**_*U*_ is given by:

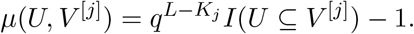

By Lemma 8, the eigenvalues of **F**^[*j*]^(**F**^[*j*]^)^*T*^ are simply one plus those calculated with Eq. S15. Since the Fourier basis diagonalizes all **F**^[*j*]^(**F**^[*j*]^)^*T*^, the eigenvalues of **FF**^*T*^ are simply the sum of those of the **F**^[*j*]^(**F**^[*j*]^)^*T*^. The eigenvalues of **FF**^*T*^ associated with the eigenvectors given by the columns of **Φ**_*U*_ are the variances of the Fourier coefficients corresponding to the epistatic interaction *U*. All of this together, we have:

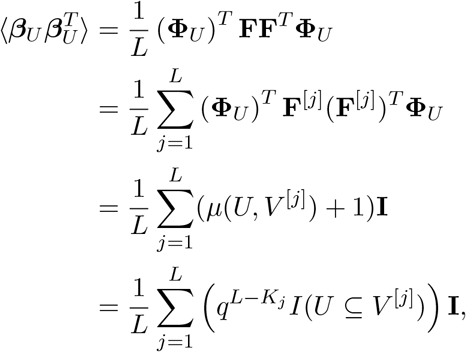

which is the desired result for the variances of the Fourier coefficients.

*Proof of Lemma 4*. Every sequence (i.e., vertex of *G*(*V*)) contains exactly one subsequence corresponding to the position indices in *V*. Therefore, each vertex is contained in exactly one edge of *G*(*V*).

*Proof of Lemma 5*. In order to prove this, we need to show (i) that the above formulation results in *L* unit normally distributed subsequence fitness values being assigned to each sequence, where each subsequence corresponds to the position indices in a neighborhood *V*^[*j*]^ (ii) that sequences share subsequence fitness values when they share the corresponding subsequence, and (iii) that the *L* subsequence fitness values are summed to produce the total fitness value assigned to each sequence.

A direct result of Definition S6.3 is that **F**^[*j*]^ has elements given by:

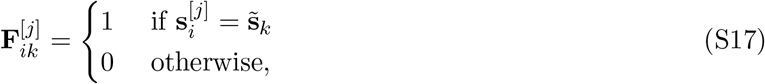

where we define 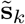 as the *k*^th^ possible subsequence of length *K_j_* (i. e., the *k*^th^ element in 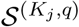. Since each hyperedge in *G*^[*j*]^ represents a subsequence of length *K_j_*, each hyperedge contains vertices that represent sequences that share subsequence fitness values in the GNK model. Therefore, letting 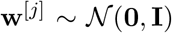 be a length *q^K_j_^* normally distributed random vector representing the subsequence fitness values randomly assigned to each subsequence, then

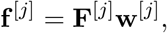

where 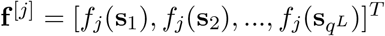 is the vector of subsequence fitness values corresponding to neighborhood *j* that are assigned to each sequence in 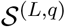. Since *G*^[*j*]^ is a 1-regular hypergraph (Lemma 4), each row of **F**^[*j*]^ contains exactly one nonzero element and therefore the subsequence fitness values of each sequence are distributed as 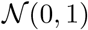, as in the original definition of the GNK model given in the main text. Additionally, the structure of the incidence matrix shown in Eq. S17 ensures that two sequences that share a subsequence corresponding to the position indices in *V*^[*j*]^ also share a subsequence fitness value in **f**^[*j*]^, as required by the GNK model.

Now, let 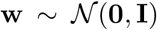 be the random vector that is the concatenation of the **w**^[*j*]^ random vectors containing subsequence fitness values. Then we have:

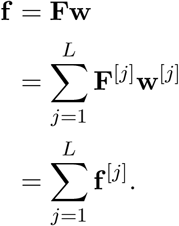

Therefore, the elements of **f** are simply the sums of the *L* subsequence fitness values corresponding to each sequence, which is the final step in definition of the GNK model given in the main text.

*Proof of Lemma 6*. This result follows immediately from recognizing that **f** = **Fw** = **Φ*β***, and therefore ***β*** = **Φ**^*T*^**Fw**. The Fourier coefficients are thus a linear transformation of a normally distributed random vector, **w**, and are therefore normally distributed with mean 〈***β***〉 = **Φ**^*T*^**F**〈**w**〉 = **0** and covariance matrix 〈***ββ***^*T*^〉 = **Φ**^*T*^ **F**〈**ww**^*T*^〉**F**^*T*^ **Φ** = **Φ**^*T*^**FF**^*T*^ **Φ**.

*Proof of Lemma 8*. Each element of **F**(*H*)**F**(*H*)^*T*^ is the inner product of two rows in **F**(*H*). Since each element in row *i* of **F**(*H*) indicates whether vertex *i* is in a particular edge, the inner product of row *i* and row *j* (*i* ≠ *j*) counts the number of edges that contain both vertex *i* and vertex *j*. Of course, this is also the number of edges connecting vertex *i* and vertex *j* in the clique multigraph, and thus the off-diagonal elements of **F**(*H*)**F**(*H*)^*T*^ are equal to the elements of **A**(*C*). The diagonal elements of **F**(*H*)**F**(*H*)^*T*^ are equal to the total number of edges containing vertex *i*, which is *L* for every vertex.

*Proof of Lemma 9*. Each vertex in a 1-regular hypergraph is in exactly one hyperedge, and thus the clique multigraph has at most one edge between any two vertices.

*Proof of Lemma 10*. First, both the lexicographic and Cartesian products result in graphs whose vertex sets are the (set) Cartesian product of the vertex sets of the multiples. Since the vertex sets of both *O*(*q*) and *K*(*q*) represent elements of the alphabet of size *q*, an *l*-fold graph product of these graphs will result in each vertex representing a sequence of length *l*.

We will prove the adjacency property of these product graphs with induction. For ease of notation, we drop the dependence of *B_l_*(*V*) on *V* and let *B_l_* ← *B_l_*(*V*). Assume that two sequences 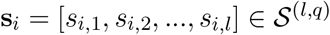 and 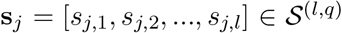 are adjacent in *B_l_* if and only if the adjacency condition, *s_i,k_* = *s_j,k_* for every *k* ∈ *V*_(*l*)_, is satisfied. We will show that these adjacency conditions remain true for *l* + 1. There are two cases to consider: (i) *l* + 1 ∈ *V* and (ii) *l* + 1 ∉ *V*.

i. (*l* + 1 ∈ *V*). Let and 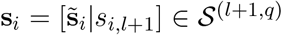 and 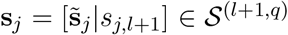 be sequences of length *l* + 1, where 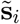 contains the first *l* elements of **s**_*i*_. Since in this case *l* + 1 ∈ *V*_(*l*+1)_, we must prove that **s**_*i*_ and **s**_*j*_ are adjacent in *B*_*l*+1_ if and only if the adjacency condition is satisfied for 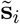 and 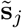 in *B_l_* (which is true by inductive assumption) and *s*_*i,l*+1_ = *s*_*j,l*+1_. By Equation S11, *B*_*l*+1_ = *B_l_□*O**(*q*) in this case. Note that the vertices in *B_l_* represent the 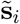 sequences of length *l* and the vertices in *O*(*q*) represent the new elements of the sequence, *s*_*i,j*+1_. According to the definition of the graph Cartesian product (Definition 1), **s**_*i*_ and **s**_*j*_ are adjacent in *B*_*l*+1_ if and only if 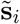 and 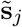 are adjacent in *B_l_* and *s*_*i,l*+1_ = *s*_*j,l*+1_. Thus, in this case, the adjacency condition remains true for *l* + 1 under the inductive assumption.
ii. (*l* + 1 ∉ *V*). In this case, *l* + 1 ∉ *V*_(*l*+1)_ and therefore we need to prove that **s**_*i*_ and **s**_*j*_ are adjacent in *B*_*l*+1_ if and only if 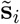 and 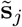 are adjacent in *B_l_* or 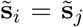. In this case, *B*_*l*+1_ = *B_l_* ∘ *K*(*q*). Due to the definition of the lexicographic product (Definition 2), **s**_*i*_ and **s**_*j*_ are adjacent in *B*_*l*+1_ if and only if (1) 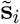 and 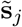 are adjacent in *B_l_* or (2) 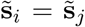 and *s*_*i,l*+1_ is adjacent to *s*_*j,l*+1_ in *K*(*q*). Since all vertices in *K*(*q*) are adjacent to one another, condition (2) simply results in **s**_*i*_ and **s**_*j*_ being adjacent in *B*_*l*+1_ if 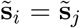. Thus, in this case, the required adjacency condition remains true for *l* + 1 under the inductive assumption.

The base case of this induction is *l* = 1. If 1 ∈ *V*, then *V*_(1)_ = {1}. Since *s*_*i*,1_ ≠ *s*_*j*,1_ for all **s**_*i*_, 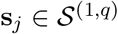 with *i* ≠ *j* (i.e., since each length-one sequence represents an element of the alphabet, none of these sequences are equal to one another), the graph *B*_1_ should contain no edges; this is indeed the case because *B*_1_ = *O*(*q*) in the case 1 ∈ *V*^[*j*]^ due to Eq. S12. Similarly, if *i* ∉ *V*^[*j*]^, then 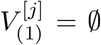 and all vertices of *B*_1_ should be adjacent to one another; this is indeed the case because *B*_1_ = *K*(*q*) when 1 ∉ *V*^[*j*]^ due to Eq. S12.

*Proof of Lemma 11*. The vertex sets of both *C*(*V*) and *B_L_*(*V*) are given by the space of sequences 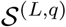. By definition, two sequences co-occur in an edge of the hypergraph *G*(*V*) if and only if they share the subsequence corresponding to the indices in *V*. Therefore two sequences are adjacent in the clique multigraph *C*(*V*) if and only if they share the subsequence corresponding to the indices in *V*. Additionally, by Lemma 9, *C*(*V*) is a simple graph. By Lemma 10, *B_L_*(*V*) is a simple graph in which two sequences are adjacent if and only if they share the subsequence corresponding to the indices in *V*. Thus, the vertex and edge sets of *C*(*V*) and *B_L_*(*V*) are equivalent, and the graphs are equivalent.

*Proof of Lemma 12*. Recall from Eq. S6 that 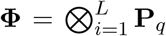, where **P**_*q*_ is a complete set of orthonormal eigenvectors of the complete graph *K*(*q*). Additionally, recognize that the adjacency matrix of the empty graph, **A**(*O*(*q*)) has every element equal to zero and therefore any nonzero vector is an eigenvector of **A**(*O*(*q*)); for our purposes we will use **P**_*q*_ as the eigenvectors of **A**(*O*(*q*)). Let the columns of **Θ**_*l*_ be orthonormal eigenvectors of the graph *B_l_*(*V*), which is defined in Lemma 10. Then we have

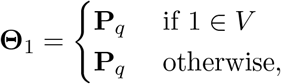

where the first and second lines on the RHS are due to **P**_*q*_ being a set of orthonormal eigenvectors of *O*(*q*) and *K*(*q*), respectively. We additionally have the recursive relation:

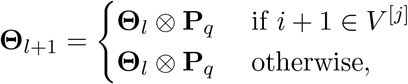

for *l* = 1, 2, … *L* – 1, where the first line on the RHS is due to Lemma 1 and the second is due to Lemma 2. Therefore, 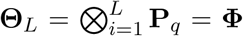. The result follows from recognizing that *C*(*V*) = *B_L_*(*V*) by Lemma 11, and therefore **Θ**_*L*_ = **Φ** are a set of orthonormal eigenvectors of **A**(*V*).

*Proof of Lemma 13*. We will prove this by induction. For ease of notation, we will drop the dependence of the **Z**_*l*_(*U*) matrices on *U*, and let **Z**_*l*_ ← **Z**_*l*_(*U*). Define **A**_*l*_ ≔ **A**(*B*_*l*_(*V*)) as the adjacency of the graph *B_l_*(*V*), which is the graph defined by Equations Eq. S11 and Eq. S12. Additionally let 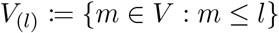 and 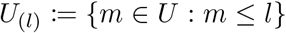 be the *l* smallest elements of *V* and *U*, respectively. Our inductive assumption will be that

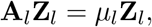

where we define 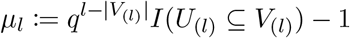. In other words, we will assume that the columns of **Z**_*l*_ are eigenvectors of **A**_*l*_ associated with the eigenvalue *μ_l_*, and then will show that **A**_*l*+1_**Z**_*l*+1_ = *μ*_*l*+1_**Z**_*l*+1_. There are four cases to consider: (i) *l* + 1 ∈ *U* and *l* + 1 ∈ *V*, (ii) *l* + 1 ∈ *U* and *l* + 1 ∉ *V*, (iii) *l* + 1 ∉ *U* and *l* + 1 ∈ *V*, and (iv) *l* + 1 ∉ *U* and *l* + 1 ∉ *V*

i. (*l* + 1 ∈ *U* and *l* + 1 ∈ *V*). In this case, *V*_(*l*+1)_ and *U*_(*l*+1)_ add the element *l* + 1 to *V*_(*l*)_ and *U*_(*l*)_, respectively. Therefore, if *U*_(*l*)_ ⊆ *V*_(*l*)_, it will be true that *U*_(*l*+1)_ ⊆ *V*_(*l*+1)_, and if 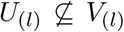, then 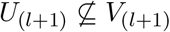. Additionally, in this case, |*V*_(*l*+1)_| = |*V*_(*l*)_| + 1, so we have

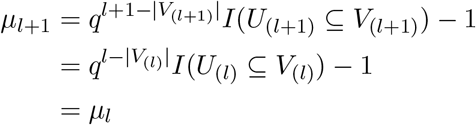 Therefore, we must show that *μ_l_* is the eigenvalue of **A**_*l*+1_ associated with the columns of **Z**_*l*+1_. In this case, 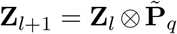. Also in this case, *B*_*l*+1_ = *B*_*l*_□*O*(*q*) (by S11), so by Eq. S1, **A**_*l*+1_ = **A**_*l*_ ⊗ **I**_*q*_. Then we have

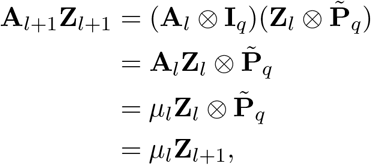

where the third line results from the inductive assumption. Thus, *μ_l_* = *μ*_*l*+1_ is the eigenvalue of **A**_*l*+1_ associated with the columns of **Z**_*l*_+1.
ii. (*l* + 1 ∈ *U* and *l* + 1 ∉ *V*). In this case, 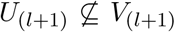 because the element *l* + 1 is in *U*_(*l*+1)_ but not *V*_(*l*+1)_. Therefore, in this case *μ*_*l*+1_ = –1, and we must prove the –1 is the eigenvalue of **A**_*l*+1_ associated with the columns of **Z**_*l*+1_. In this case, 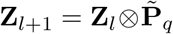. Also, in this case, *B*_*l*+1_ = *B_l_*∘*K*(*q*), so by Eq. S2,

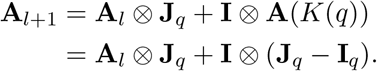 Then we have,

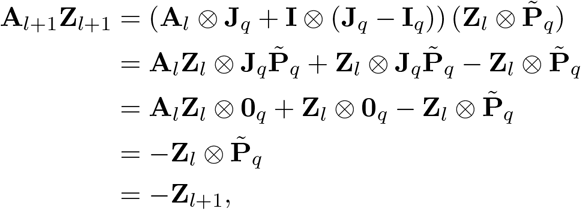

where the third line results from recognizing that each column of 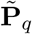 sums to zero, so 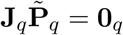, where **0**_*q*_ is the *q* × *q* matrix of all zeros. Thus, *μ*_*l*+1_ = –1 is the eigenvalue of **A**_*l*+1_ associated with the columns of **Z**_*l*_+1.
iii. (*l* + 1 ∉ *U* and *l* + 1 ∈ *V*). In this case, the element *l* + 1 is in *V*_(*l*+1)_ but not *U*_(*l*+1)_. Therefore, if *U*_(*l*)_ ⊆ *V*_(*l*)_, it will be true that *U*_(*l*+1)_ ⊆ *V*_(*l*+1)_, and if 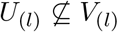, then 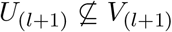. Additionally, in this case, |*V*_(*l*+1)_| = |*V*_(*l*)_| + 1, so, as in case (i), we have *μ*_*l*+1_ = *μ_l_*, and we must prove the *μ_l_* is the eigenvalue of **A**_*l*+1_ associated with the columns of **Z**_*l*+1_. In this case, **Z**_*l*+1_ = **Z**_*l*_ ⊗ **1**_*q*_ and **A**_*l*+1_ = **A**_*l*_ ⊗ **I**_*q*_ (as in case (i)). Then,

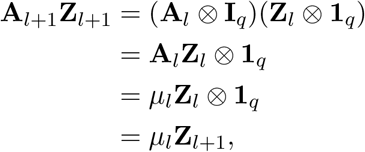

where the third line results from the inductive assumption. Thus, *μ_l_* = *μ*_*l*+1_ is the eigenvalue of **A**_*l*+1_ associated with the columns of **Z**_*l*+1_.
iv. (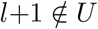 and 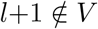). In this case, *V*(*l*_(*l*+1)_ = *V*(*l*) and *U*_(*l*+1)_ = *U*_(*l*)_, so 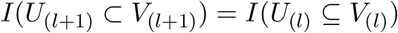 and 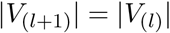. Therefore,

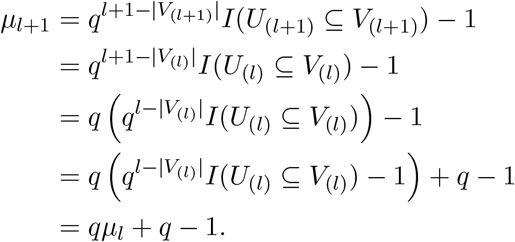 Thus, we must prove that *qμ_l_* + *q* – 1 is the eigenvalue of **A**_*l*+1_ associated with the columns of **Z**_*l*+1_. In this case, **Z**_*l*+1_ = **Z**_*l*_ ⊗ **1**_*q*_ (as in case (iii)) and **A**_*l*+1_ = **A**_*l*_ ⊗ **J**_*q*_ + **I** ⊗ (**J**_*q*_ – **I**_*q*_) (as in case (ii)). Then, we have

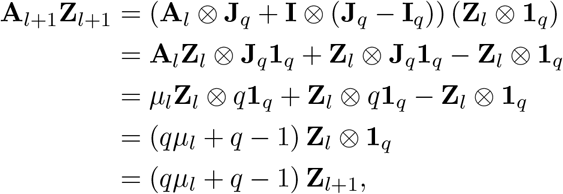

where the third line results from the inductive assumption and recognizing that **J**_*q*_ **1**_*q*_ = *q***1**_*q*_. Thus, *μ*_*l*+1_ = (*qμ_l_* + *q* – 1) is the eigenvalue of **A**_*l*+1_ associated with the columns of **Z**_*l*+1_.

We additionally have four analogous base cases for the induction: (i) 1 ∈ *U* and 1 ∈ *V*, (ii) 1 ∈ *U* and 1 ∉ *V*, (iii) 1 ∉ *U* and 1 ∉ *V*, and (iv) 1 ∉ *U* and 1 ∉ V:

i. (1 ∈ *U* and 1 ∈ *V*). In this case, *U*_(1)_ = *V*_(1)_ = {1}, so *I*(*U*_(1)_ ⊆ *V*_(1)_) = 1, |*V*_(1)_| = 1, and therefore *μ*_1_ = 0. Additionally, **A**_1_ = **A**(*O*(*q*)) = **0**_*q*_, so **A**_1_**Z**_1_ = *μ*_1_**Z**_1_ = **0**_*q*_.
ii. (1 ∈ *U* and 1 ∉ *V*). In this case, 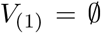, so 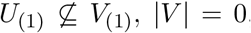, and *μ*_1_ = –1. Additionally, 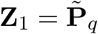 and **A**_1_ = **A**(*K*(*q*)) = **J**_*q*_ – **I**_*q*_, so we have 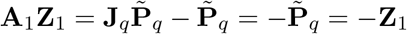.
iii. (1 ∉ *U* and 1 ∈ *V*). In this case 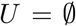 and *V* = {1}, so *U*_(1)_ ⊆ *V*_(1)_, |*V*| = 1, and *μ*_1_ = 0. Since **A**_1_ = **A**(*O*(*q*)) = **0**_*q*_, then **A**_1_**Z**_1_ = *μ*_1_**Z**_1_ = **0**.
iv. (1 ∉ *U* and 1 ∉ *V*). In this case, 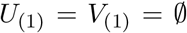, so *U*_(1)_ ⊆ *V*_(1)_, | *V*_(1)_ | = 0 and thus *μ*_1_ = *q* – 1. Additionally, **Z**_1_ = **1**_*q*_ and **A**_1_ = **A**(*K*(*q*)) = **J**_*q*_ – **I**_*q*_, so we have

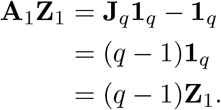

Thus, *μ*_1_ is the eigenvalue of **A**_1_ associated with the columns **Z**_1_ in each of these base cases.

By this induction, we have proved that *μ_L_* is the eigenvalue of **A**_*L*_ associated with the columns of **Z**_*L*_. It is clear to see that *μ_L_* = *μ*(*U, V*). The result then follows from recognizing that, due to Lemma 11, *B_L_*(*V*) = *C*(*V*) and therefore **A**_*L*_ = **A**(*V*). Thus, from the induction, the columns of **Z***_L_*(*U*) are eigenvectors *A*(*V*) associated with the eigenvalue *μ_L_* = *μ*(*U, V*).

*Proof of Lemma 14*. For this proof, recall that the *i*^th^ row of 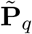 encodes the *i*^th^ element of the alphabet. We will denote each of these encodings as **p**_*q*_(*s*), where *s* is an element of the alphabet (i.e., each **p**_*q*_(*s*) is a row of 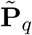). Let 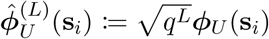 for *i* = 1, 2, …*q^L^* be the unnormalized rows of **Φ**_*U*_. These can be defined recursively. In particular, we have

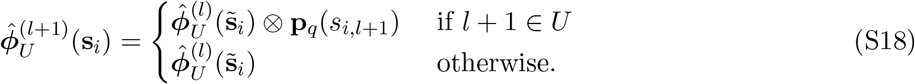

for *l* = 1, 2, …, *L*, where 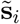 are the first *l* positions of **s**_*i*_,

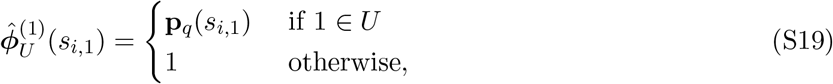

and 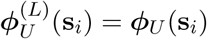. We can then recursively define the **Φ**_*U*_ matrix, by letting

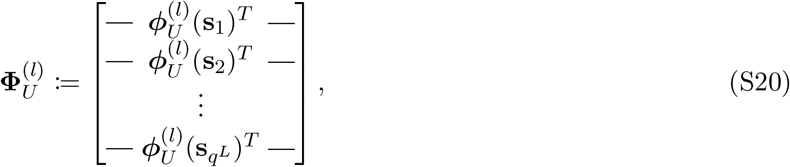

For a given 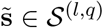, there are *q* sequences in 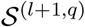 whose first *l* positions are 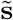. Further, each of these sequences has a unique element in the final position. Thus,

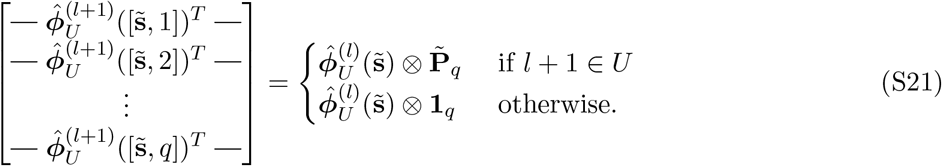

Now let, 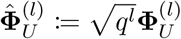. Applying Eq. S21 to each row in 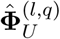 results in:

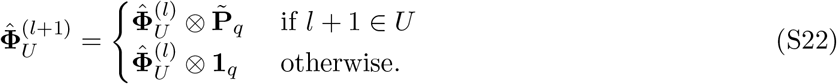

which is equivalent to the recursion in Eq. S13 that defines **Z**_*l*_(*U*). Additionally, repeated application of Eq. S19 to each element in the alphabet results in the equivalent base case to Eq. S14. Carrying out the recursion of Eq. S21 to *l* = *L* then gives

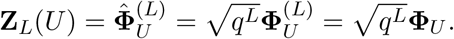

### S6.4 The sparsity of GNK fitness functions

Here we prove our main result regarding the sparsity of the Fourier coefficients of fitness functions sampled from the GNK model, which is summarized in Eq. 6. First we re-state this result formally.

#### Theorem 2.

*Let S*(*f*) ≔ #supp(***β***) *be the sparsity of a fitness function f of sequences of length L and alphabet size q with Fourier coefficients ***β***, where supp(***β***) is the set of non-zero elements of ***β*** and # is the counting measure. Then for any* 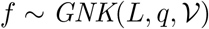,

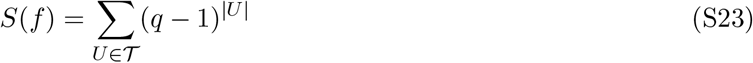

*almost surely, where 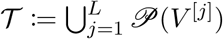 is the union of the powerset of each neighborhoods*.

*Proof of Theorem 2*. Theorem 1 shows that all the Fourier coefficients associated with an epistatic interaction *U* are deterministically zero if *U* ⊈ *V*^[*j*]^ for *j* = 1, 2, …, *L*, which can be alternatively stated as 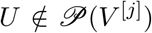 for *j* = 1, 2, …, *L*. Recalling that 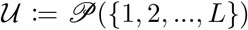, the epistatic interactions with nonzero Fourier coefficients are the 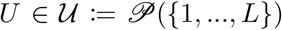 for which there exists a *j* ∈ 1, 2, …, *L* such that 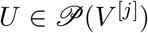. Since 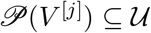, we have

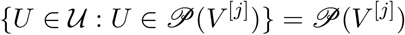

and further,

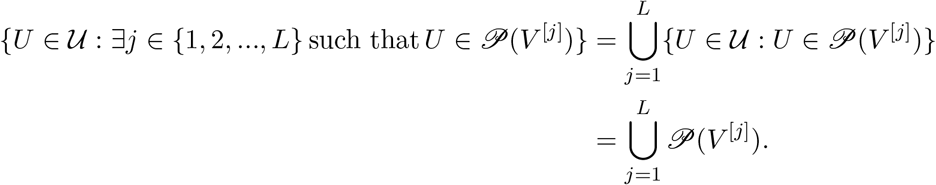

There are (*q* – 1)^|*U*|^ Fourier coefficients associated with each 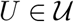, so letting 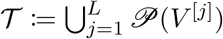, we have

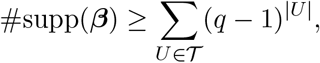

where the bound results from recognizing that the RHS sums overs all Fourier coefficients that are *deterministically* zero, but the coefficients with nonzero variances may still equal zero. However, recognizing that each *β* ∈ ***β*** with nonzero variance is a normal random variable that can equal zero with zero probability, we have

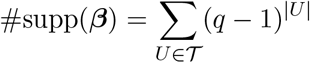

almost surely.

### S6.5 The sparsity of GNK fitness functions with standard neighborhood schemes

Here we prove our results regarding the sparsity of GNK fitness functions with standard neighborhood schemes. In particular, we prove (i) an upper bound on the sparsity of any GNK fitness function with constant neighborhood sizes (Eq. in the main text), (ii) the sparsity of GNK fitness functions with Block Neighborhoods (Eq. 12), (iii) the sparsity of GNK fitness functions with Adjacent Neighborhoods (Eq. 13) and (iv) the expected sparsity of GNK fitness functions with Random Neighborhoods. Below we restate each of these results formally, and provide proofs. We start with the bound of Eq..

#### Proposition 3.

*Let 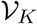 be a set of neighborhoods where K_j_* = *K for j* = 1, 2, …, *L and* 1 ≤ *K* ≤ *L. Then, the sparsity of any 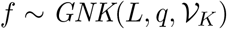 is bounded above by*:

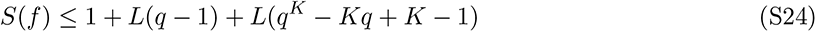

*Proof*. Let 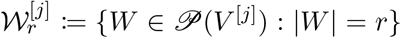 be the number of elements of the powerset of neighborhood *j* with cardinality *r*. Additionally define

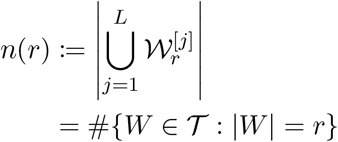

as the number of elements in the union of powersets with cardinality *r*. For any set of neighborhoods, we have *n*(0) = 1 and *n*(1) = *L*. Additionally, for any 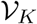 with constant neighborhood size *K*, *n*(*r*) = 0 for *r* > *K*. Then, for *r* = 2, 3…, *K*, we have

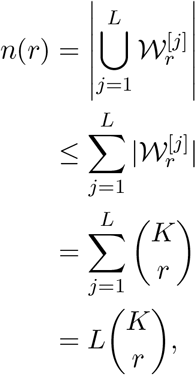

where the second line results from the union bound and the third from recognizing that there are 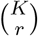 sets of cardinality *r* in the powerset of a set with *K* elements. Using this within Theorem 2 we then have

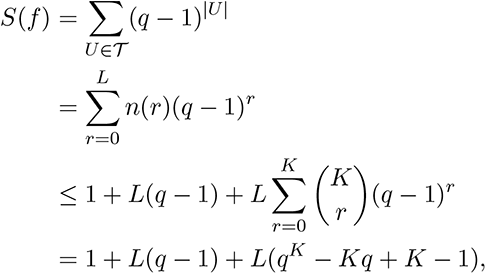

where the final line results from the binomial theorem.

The next result calculates the sparsity of GNK fitness functions with Block Neighborhoods.

#### Proposition 4.

*Given an L, q, and K satisfying L* mod *K* = 0 (*i. e., K must be set such that L is a multiple of K), define a Block Neighborhood as*

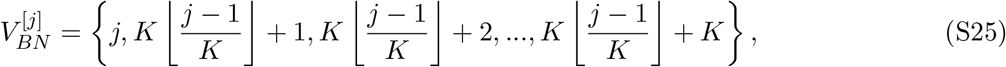

*where* ⌊⌋ *is the floor operator, and we assume, without loss of generality, that the positions in each block are adjacent. Further let* 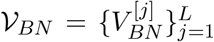 *be a set of L Block Neighborhoods. Then, the sparsity of a fitness function f sampled from 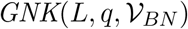 is given by*:

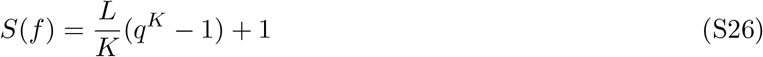

*Proof*. There are 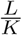 blocks. The blocks are fully connected, so all 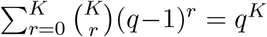 Fourier coefficients corresponding to intra-block epistatic interactions are nonzero. The only epistatic interaction shared by the blocks is the zeroth order interaction, so each block contributes (*q^K^* – 1) unique nonzero Fourier coefficients, and the total number of nonzero Fourier coefficients is given by Eq. S26, where the final addition of one is due to the shared zeroth order interaction.

Similarly, the sparsity of GNK fitness with Adjacent Neighborhoods is shown in the following proposition.

#### Proposition 5.

*Given an L, q, and 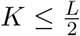, define an Adjacent Neighborhood as*

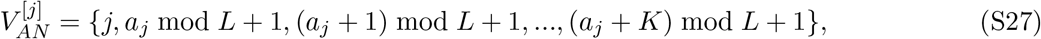

*where we define* 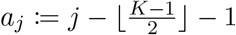. *Further let* 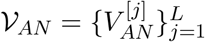 *be a set of L Adjacent Neighborhoods. Then, the sparsity of any* 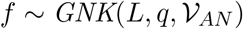 *is given by*:

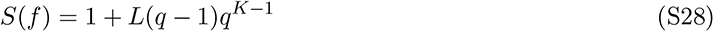

*Proof*. Define 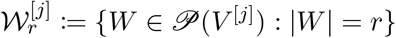 and

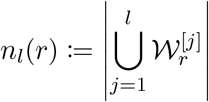

for *l* = 1, 2, …, *L*. For *l* ≤ *L* – *K* + 1, and *r* = 1, 2, …, *K*, we have

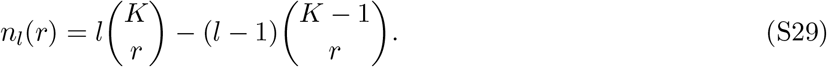

This can be shown by induction. In particular, assume Eq. S29 is correct for *l* < *L* – *K* + 1 and then we find:

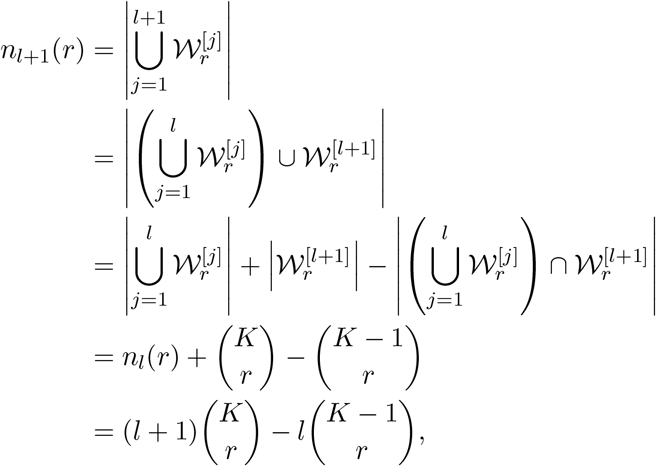

where the fourth line results from recognizing that *V*^[*l*+1]^ when *l* + 1 ≤ *L* – *K* + 1 contains exactly one position that is not in 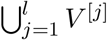; there are then 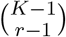 sets in 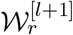 that contain this element and are thus unique to 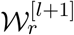, which leads to

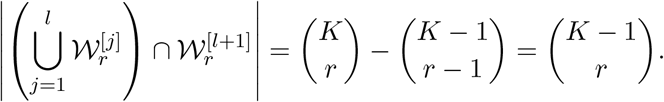

It is clear that 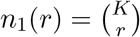, and thus Eq. S29 is proved by induction for *l* ≤ *L* – *K* + 1.

Eq. S29 accounts for redundancies in 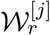 that result from overlapping positions in the neighborhoods, without considering periodicity. There are additional redundancies that occur when *l* > *L* – *K* + 1 due to the periodicity of the neighborhoods. In particular, for *l* = *L* – *K* + 2, …, *L*, due to periodicity *V*^[*l*]^ contains (*l* + *k*) mod *L* – 1 additional positions that are already in 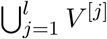 (outside of those that are already 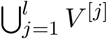 due to non-periodic overlap). Therefore 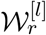 contains 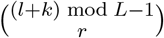 additional sets that are already in 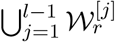 due to periodicity. Then we have, for *l* = *L* – *K* + 2, …, *L*:

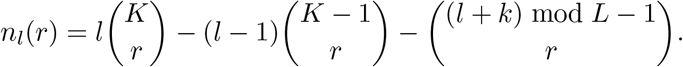

At *l* = *L*, we then have

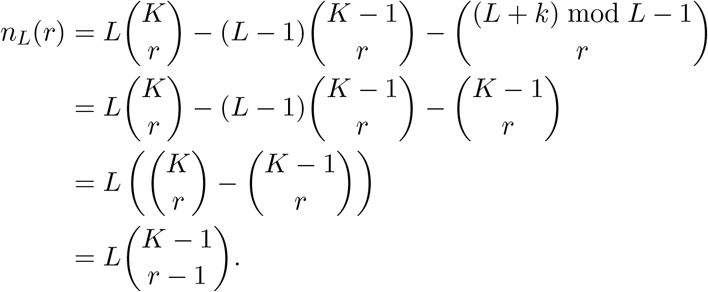

The result follows from recognizing that *n_L_*(0) = 1, and therefore

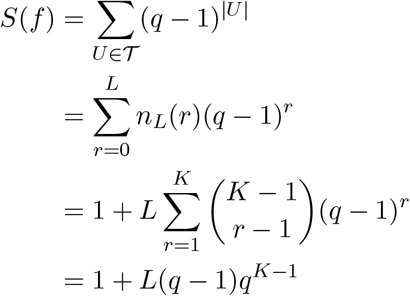

where the final line is due to the binomial theorem.

Additionally, we are able to calculate the expected sparsity of GNK fitness functions with Random Neighborhoods, which is shown in the following result. The proof of this follows the analogous calculations of ref. 31, and we correct a mistake in their calculations.

#### Proposition 6.

*Let* 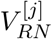 *be a set of cardinality K, where the first K* – 1 *elements are selected uniformly at random from* {1, 2, …, *L*} *without replacement, and the final element is *j*. Let* 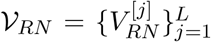 *be a collection of such sets. Then, the expected sparsity of a fitness function f sampled from* 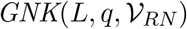, *with the expectation taken over the possible realizations of* 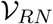, *is given by*:

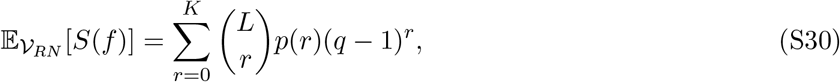

*where*

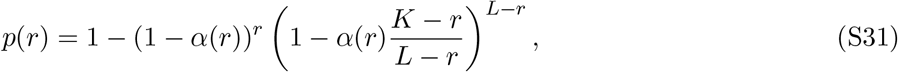

*and* 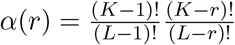

*Proof*. Consider a set *W* ⊆ {1, 2, …, *L*} of cardinality *r*. Define *α*(*r*) as the the probability that *W* is a subset of the random neighborhood *V*^[*j*]^ given that *j* ∈ *W*, which is given by

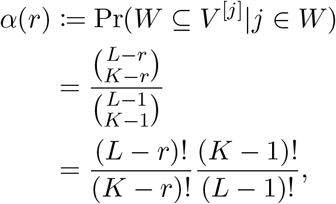

where 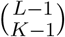 is the total number of ways to construct *V*^[*j*]^ and 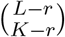 is the number of ways to construct *V*^[*j*]^ such that every element of *W* is in *V*^[*j*]^. The probability that *W* is a subset of the random neighborhood *V*^[*j*]^ given that *j* ∉ *W* is similarly given by:

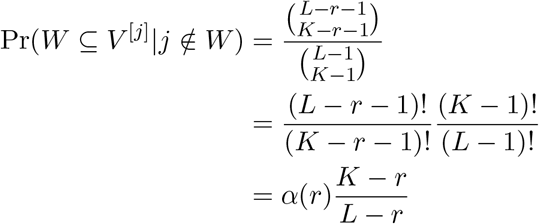

There are *r* neighborhoods *V*^[*j*]^ for which *j* ∉ *W*, and *L* – *r* neighborhoods for which *j* ∈ *W*. Define *p*(*r*) as the probability that *W* is a subset of at least one, which is then:

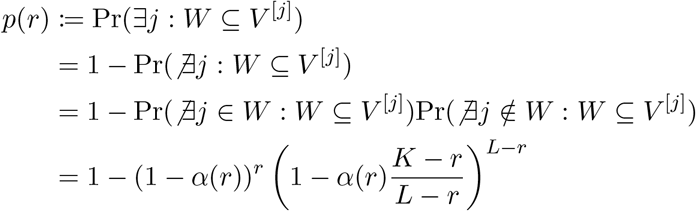

There are 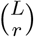 sets of cardinality *r*; in expectation 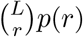 will be subsets of at least one neighborhood, and will therefore represent epistatic interactions or order *r* corresponding to (*q* – 1)^*r*^ nonzero Fourier coefficients. Eq. S30 follows from summing over all possible cardinalities *r*.

## S7 Extension to non-constant alphabet sizes

It is common that alphabet sizes are not constant at every position. Here we generalize our formal results to the case of “hybrid alphabets”, where alphabet sizes may differ at each position. Consider the case where the alphabet size at each position is given by the length *L* vector **q** = [*q*_1_, *q*_2_, …, *q_L_*] and let 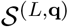 be the space of all sequences corresponding to the alphabet sizes in **q**. Denote as *H*(*L*, **q**) the Generalized Hamming graph whose vertex set is 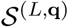 and whose edges connect sequences that differ in exactly one position [56]. The Generalized Hamming graph can be constructed as an *L*-fold graph Cartesian product:

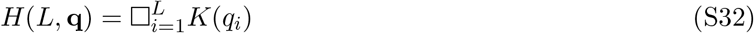

We then have the following result for the Fourier basis corresponding to these Generalized Hamming graphs, which follows straightforwardly applying Lemma 3 to Equation Eq. S32.

### Proposition 7.

*An orthonormal set of eigenvectors of the Graph Laplacian of the Generalized Hamming graph H*(*L*, **q**) *is given by*

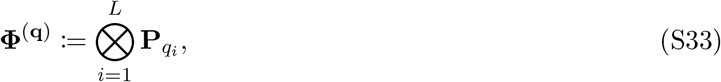

*where* **P**_*qi*_ *is defined in* Eq. S5.

In this basis, each epistatic interaction *U* is represented by ∏_*k∈U*_(*q_k_* – 1) columns of **Φ**^(*q*)^.

The Fourier basis of Eq. S33 can be used to represent fitness functions of sequences with non-constant alphabet sizes, **q**. The GNK model can also be defined analogously to the definition given in the Materials and Methods section for the case of non-constant alphabet sizes; in particular let 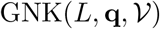 be the distribution over fitness functions of sequences of length *L* with non-constant alphabet sizes given by **q** and neighborhood set 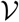. We then have the following results regarding the distribution and support of the Fourier coefficients of fitness functions sampled from this distribution. We present these results without proof, though it is straightforward to see how to adapt the proof of Theorems 1 and 2 to prove these.

### Theorem 3.

*Let* 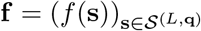 *be the complete vector of evaluations of a fitness function* 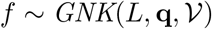. *Then the Fourier coefficients of f, **β*** = (**Φ**^(*q*)^)^*T*^**f**, *are distributed according to 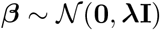. Let ***β***_U_ be the length ∏_k∈U_* (*q_k_* – 1) *sub-vector of ***β*** representing the epistatic interaction U. Then the variance of every element of ***β***_U_ is given by*:

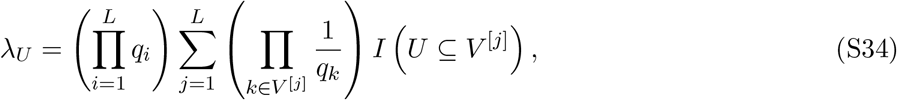

*where I* (*U* ⊆ *V*^[*j*]^) *is an indicator function that is equal to one if U is a subset of or equal to V*^[*j*]^ *and zero otherwise*.

### Theorem 4.

*The sparsity of any* 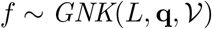 *is given by*

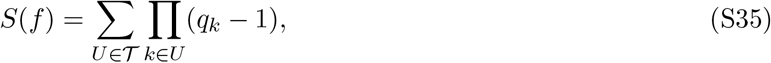

*almost surely*.

## Footnotes

1 In the original definition of the model, *N* is used for the sequence length, but here we reserve *N* for the number of observed measurements.

2 In a quirk of common terminology, a signal is called “sparse” when it contains many zero coefficients, but the “sparsity” is formally defined as the number of nonzero coefficients. Thus, a ‘sparse’ signal has *low* “sparsity”.

## Notes

### Competing Interest Statement

Jennifer Listgarten is on the Scientific Advisory Board for Foresite Labs and Patch Biosciences.

